# Probabilistic Inference on Virtual Brain Models of Disorders

**DOI:** 10.1101/2024.02.21.581243

**Authors:** Meysam Hashemi, Abolfazl Ziaeemehr, Marmaduke M. Woodman, Spase Petkoski, Viktor K. Jirsa

## Abstract

Connectome-based models, also known as Virtual Brain Models (VBMs), have been well established in network neuroscience to investigate pathophysiological causes underlying a large range of brain diseases. The integration of an individual’s brain imaging data in VBMs has improved patient-specific predictivity, although Bayesian estimation of spatially distributed parameters remains challenging even with state-of-the-art Monte Carlo sampling. VBMs imply latent nonlinear state space models driven by noise and network input, necessitating advanced probabilistic machine learning techniques for widely applicable Bayesian estimation. Here we present Simulation-Based Inference on Virtual Brain Models (SBI-VBMs), and demonstrate that training deep neural networks on both spatio-temporal and functional features allows for accurate estimation of generative parameters in brain disorders. The systematic use of brain stimulation provides an effective remedy for the non-identifiability issue in estimating the degradation of intra-hemispheric connections. By prioritizing model structure over data, we show that the hierarchical structure in SBI-VBMs renders the inference more effective, precise and biologically plausible. This approach could broadly advance precision medicine by enabling fast and reliable prediction of patient-specific brain disorders.

## 1. Introduction

Model-based inference involves constructing a statistical or mechanistic model that captures the essential features of the data-generating process and the underlying structure or relationships in the observed data (Anderson, 2008). It emphasizes the incorporation of domain knowledge (Gelman et al., 2013), hypothesis testing (MacKay, 2003), interpretability (Hastie et al., 2009), generalizability (Vapnik, 1999), and causality (Pearl, 2009). It also often provides superior performance over purely data-driven methods, particularly in the context of brain disorders, such as epilepsy (Hashemi et al., 2020; Wang et al., 2023; Jirsa et al., 2023), and Alzheimer’s disease (Triebkorn et al., 2022; Yalcinkaya et al., 2023). One class of computational network models commonly used to analyze functional neuroimaging modalities, such as fMRI, MEG, and EEG, is the class of connectome-based models (Ghosh et al., 2008; Sanz-Leon et al., 2015; Bassett and Sporns, 2017), also known as Virtual Brain Models (VBMs; Sanz Leon et al. (2013); Jirsa et al. (2023)). Each node in these models corresponds to a brain region (Bullmore and Sporns, 2009; Sporns, 2016). The collective behavior of these nodes over time is then described using a dynamical model of average neuronal activity, known as neural mass models (Deco et al., 2008, 2011; Breakspear, 2017). The connectome-based models incorporate individual structural brain imaging data (Melozzi et al., 2019; Hashemi et al., 2021), typically diffusion-weighted imaging data, to estimate the edge weights (Hagmann et al., 2007; Van Essen et al., 2012; Schirner et al., 2015, 2023). The structural connectivity imposes a constraint on the brain dynamics, allowing for the personalized simulation of the brain’s (ab)normal activities and their associated imaging recordings, potentially informing targeted interventions (Proix et al., 2017; Olmi et al., 2019; Wang et al., 2023; Jirsa et al., 2023).

During aging or some diseases, the SC is reported to deteriorate (Antonenko and Flöel, 2013; Damoiseaux, 2017; Zuo et al., 2017), or be perturbed (Stam, 2014; Ozdemir et al., 2020; Menardi et al., 2021), particularly with respect to the number of inter- and intra-hemispheric fibers within tracts and fiber density (Puxeddu et al., 2020; Petkoski et al., 2023; Lavanga et al., 2023). These observations, however, do not map trivially on functional data, such as fMRI (Stumme et al., 2022; Jockwitz et al., 2023; Krämer et al., 2023). This could be due to several factors, notably those related to the structure of the generative model family: these features include a degenerate mapping between Structural Connectivity (SC) and Functional Connectivity (FC), high dimensionality of the parameter space, and degeneracy induced by network effects in the latent state space, all introducing potential non-identifiability. These sources of indeterminacy stymie even state-of-the-art methods, which invites us to find ways to improve identifiability and employ algorithms better equipped for dealing with degeneracy.

State-of-the-art methods for inverting models of neuroimaging data vary in realism and complexity of the resulting estimate. Methods such as Dynamical Causal Modeling (Friston et al., 2003, 2014) combine sophisticated statistical models and fixed variational schemes to address specific mechanistic hypotheses in neuroimaging datasets but scale poorly in terms of the size of the hypothesized network, and its mean-field variant ignores - by definition-the correlation between parameters. The Digital Twin Brain (Lu et al., 2022), on the other hand, models a very large network of spiking neurons and employs a Kalman filter scheme (a type of recursive filtering algorithm) to fit the data, yet it remains difficult to assess both the realism of the network connectivity and the interpretability of the model parameters. The VBMs aim to push the frontier forward by combining scalability to realistically detailed whole-brain models with recent probabilistic inference schemes to yield reliable, mechanistically interpretable estimates (Hashemi et al., 2020, 2021, 2023). The resulting improvements in the state-of-the-art are achieved through parameterizations, which decorrelate model parameters (e.g., to take advantage of parallel simulations). The resulting models scale well with data resources while remaining tractable for probabilistic inference on personalized whole-brain functional data. This work highlights the importance of exploring flexible and scalable alternative methods that may offer unique advantages in terms of realism, interpretability, and uncertainty quantification through Bayesian estimation for better downstream decision-making processes.

Using random simulations, Markov Chain Monte Carlo (MCMC) is the gold standard technique for carrying out the Bayesian inference (Andrieu et al., 2003; Murphy, 2022; McElreath, 2020). MCMC is a powerful class of computational algorithms for sampling from a probability distribution, in which the sampling process does not require knowledge of the whole distribution. In Bayesian inference, MCMC methods are often used for unbiased sampling from distributions, which is asymptotically exact in the limit of infinite runs, but require explicit evaluation of the likelihood function (Hashemi et al., 2020, 2021). The Markovian (sequential) structure of MCMC methods interacts poorly with the highly multimodal, non-convex posteriors of generative VBMs, where nonlinear latent dynamics generally imply multistability in a high dimensional state space. Hence, MCMC methods require restrictive reparametrizations to deal with geometrical issues (Betancourt et al., 2014; Betancourt, 2016), immense computational cost (Baldy et al., 2024), or intricately designed sampling strategies (Hashemi et al., 2020; Jha et al., 2022) to efficiently converge (Gabrié et al., 2022; Baldy et al., 2023).

Simulation-Based Inference (SBI; Cranmer et al. (2020); Boelts et al. (2022)) or likelihood-free inference (Papamakarios et al., 2019b; Brehmer et al., 2020) sidesteps problems with posterior geometry and algorithm sequentiality entirely: SBI uses the generative model to map samples from the prior to a corresponding set of data features, and then it takes a maximum likelihood estimate of a Bayesian regression of model parameters on data features. In practice, to ensure the resulting approximate posterior density is sufficiently expressive, these methods employ deep neural networks to parametrize or construct directly the approximated density (Greenberg et al., 2019; Gonçalves et al., 2020; Hashemi et al., 2023). In these deep network approaches, a simple base probability distribution (i.e., prior) is transformed into a more complex distribution (i.e., posterior) through a sequence of invertible transformations (Rezende and Mohamed, 2015; Papamakarios et al., 2019a)). In particular, the advanced machine learning techniques based on unsupervised generative models offer efficient reconstruction of the probability density functions and highly expressive transformations with low-cost computation (Kobyzev et al., 2020; Papamakarios et al., 2021), hence, efficient Bayesian inference on complex high-dimensional systems. SBI implies computationally intensive simulation and training stages, but performed only once, and subsequent inference is extremely efficient as it requires only a forward pass of the neural network on a vector of data features to construct a posterior distribution (in order of seconds, due to amortized strategy; Hashemi et al. (2023)).

In this study, we use an exact mean-field model of spiking neurons at each parcellated brain region, displaying a variety of dynamics such as multi-stability, in which the emergent dynamics are constrained by personalized anatomical data (SC matrix). We aim to reliably estimate the full posterior distribution of control parameters in VBMs, by training deep neural networks on a low-dimensional representation of fMRI BOLD data, such as functional connectivity (FC; Friston (1994); Honey et al. (2009); Greicius et al. (2003)), and Functional Connectivity Dynamics (FCD; Zalesky et al. (2014); Hansen et al. (2015); Lurie et al. (2020)). We use the term *spatio-temporal data features* to refer to the statistical moments derived from BOLD time-series, while we refer to the summary statistics extracted from FC and FCD matrices as *functional data features*. The main motivation for using the SBI approach is to leverage the advantages of fast and parallel simulations for efficient and flexible inference using data features, accompanied by uncertainty quantification rather than a single point estimation (Wang et al., 2019; Kong et al., 2021).

Note that opting for SBI over MCMC sampling will not offer a straightforward solution to the inference challenges at the whole-brain level. A generative model defines the joint probability distribution of both observed data and latent variables or parameters of the model, enabling the generation of synthetic data samples. Hence, the performance of inference relies on both the level of information contained in the observed data, and the efficiency of model structure to consistently recognize that data. Accordingly, to address non-identifiability issues using generative models such as VBMs, we take the following approaches: (i) Enhancing the inference by incorporating more information, e.g., through data augmentation; (ii) Increasing the probability transition between states in latent dynamics, e.g., through intervention by stimulation; (iii) Restructuring the model configuration space, e.g., through hierarchical reparameterization, to facilitate more efficient exploration of the posterior distribution.

Our results on the uncertainty quantification using SBI show that FCD is more informative than FC for inference on perturbed connections, but using both data features provides stronger model evidence against the fitted data. Nevertheless, correlations between different brain regions and the associated dynamics (functional data features) do not provide sufficient information for inferring heterogeneous excitability at whole-brain level. We demonstrate that training deep neural density estimators (Papamakarios et al., 2019a; Kobyzev et al., 2020) by including the spatio-temporal data features provides more accurate inference on generative parameters. As an alternative to such data augmentation, the perturbation in brain dynamics by stimulation can provide an efficient remedy for the non-identifiability issue in the estimation of degradation in intra-hemispheric connections (within the limbic system). By prioritizing model structure over data, we show that the hierarchical structure in VBMs proves to be more effective than pooled (homogeneous) and unpooled (heterogeneous) modeling, to infer the control parameters from functional data. This set of control parameters comprises the excitability across brain regions, a global scaling factor on the connectome, and the level of (inter- and intra-hemispheric) degradation in the connectome. The SBI-VBMs approach is now available on the cloud platform EBRAINS (Schirner et al., 2022) to assist users in uncovering more realistic brain dynamics that underlie brain diseases, within a Bayesian causal framework.

## 2. Materials and methods

### 2.1 Structural connectivity

The Virtual Brain Models (VBMs) emphasize the structural network characteristics of the brain by representing the brain regions as nodes, which are connected via a structural connectivity matrix representing white matter fibre tracts (Sanz-Leon et al., 2015). The Structural Connectivity, *SC* ∈ ℝ ^*N×N*^ with *N* parcelled regions, was derived from a probabilistic tractography-based connectome using generally available neuroimaging software, such as The Virtual Brain (TVB; Sanz Leon et al. (2013); Schirner et al. (2015)). The T1-weighted MRI images were processed to obtain the brain parcellation, whereas Diffusion-weighted (DW-MRI) images were used for tractography. With the generated fiber tracts and with the regions defined by the brain parcellation, the connectome was built by counting the fibers connecting all regions. Using Desikan-Killiany parcellation (Desikan et al., 2006) in the reconstruction pipeline, the patient’s brain was divided into *N* = 88 cortical and subcortical regions. The connectome was normalized so that the maximum value is equal to one.

### 2.2 The virtual brain models

To ensure a realistic in-silico experiment and therefore assess the function-structure link during the brain disorders via VBMs, we simulated the blood-oxygen level dependent (BOLD) fMRI using a whole-brain network model that couples exact mean-field representation of spiking neurons through the weighted edges in the SC matrix. Assuming a Lorentzian distribution on the membrane potentials across decoupled brain regions, with half-width Δ centered at η, the macroscopic dynamics associated with a local network node is governed by a neural mass model (NMM) derived analytically as the limit of infinitely all-to-all coupled quadratic integrate-and-fire (QIF) neurons (Montbrió et al., 2015):

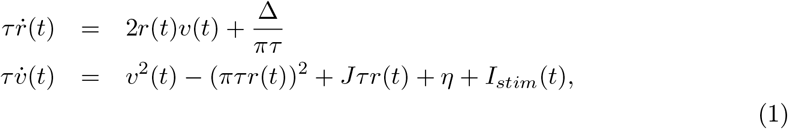

where *v* and *r* are the average membrane potential and firing rate, respectively, at each brain region. The excitability *η* = *−*4.6, the synaptic weight *J* = 14.5, the spread of the heterogeneous noise distribution Δ = 0.7, and the characteristic time constant *τ* = 1, are tuned so that each decoupled node is in a bistable regime, exhibiting a down-state stable fixed point and an up-state stable focus in the 2D phase space (Montbrió et al., 2015).

The bistability is a fundamental property of regional brain dynamics to ensure a switching behavior in the data (e.g., to generate FCD), that has been recognized as representative of realistic dynamics observed empirically (Rabuffo et al., 2021). The input current I_stim_ represents the stimulation to selected brain regions, which increase the basin of attraction of the up-state in comparison to the down-state, while the fixed points move farther apart (Rabuffo et al., 2021).

By coupling the brain regions via an additive current in the average membrane potential equations, the dynamics of the whole-brain network can be described as follows (Rabuffo et al., 2021; Fousek et al., 2022):

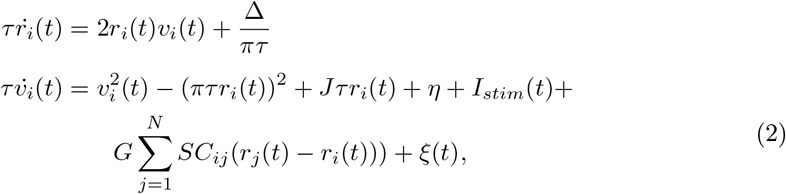

where *G* is the network scaling parameter that modulates the overall impact of brain connectivity on the state dynamics (see Supplementary, Fig S1), *SC*_*i,j*_ denotes the symmetrical connection weight between *i*_*th*_ and *j*_*th*_ nodes with *i, j* ∈ *{*1, 2, …, N *}*, and the dynamical noise ξ(*t*) *∼ 𝒩* (0, σ^2^) follows a Gaussian distribution with mean zero and variance *σ*^2^ = 0.03. The brain network model was implemented in TVB software (Sanz Leon et al., 2013; Rabuffo et al., 2021) and equipped with BOLD forward solution comprising the Balloon-Windkessel model (Stephan et al., 2007) applied to the firing rate. In addition, we have implemented the model in C++, which is around 2 times faster than our Python implementation with Just-in-Time (JIT) compilation, and also CUDA-based GPU, which provides up to 100 times faster computational cost through parallelization (see Supplementary, Fig S2).

The approach of virtually modeling the brain connectivity in disorders provides the basis to investigate whether a specific modification in connectome affects the observed brain function in a causal sense. We specifically tested whether the disorder is related to interhemispheric and intra-hemispheric connections in SC, by applying two spatial masks on the subject-specific connectome (normalized by the scaling factor G) as follows:

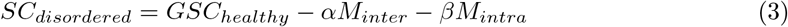

where parameters *α* and *β* are the unknown normalized intensity of degradation in SC (as the target of parameter estimation in the range [0-1]), according to the spatial masks *M*_*inter*_ and *M*_*intra*_, on inter-hemispheric and intra-hemispheric connections, respectively. In this study, the M_intra_ mask indicate the afferent and efferent connections originating from the limbic system (see Fig 1).

**Fig. 1.**
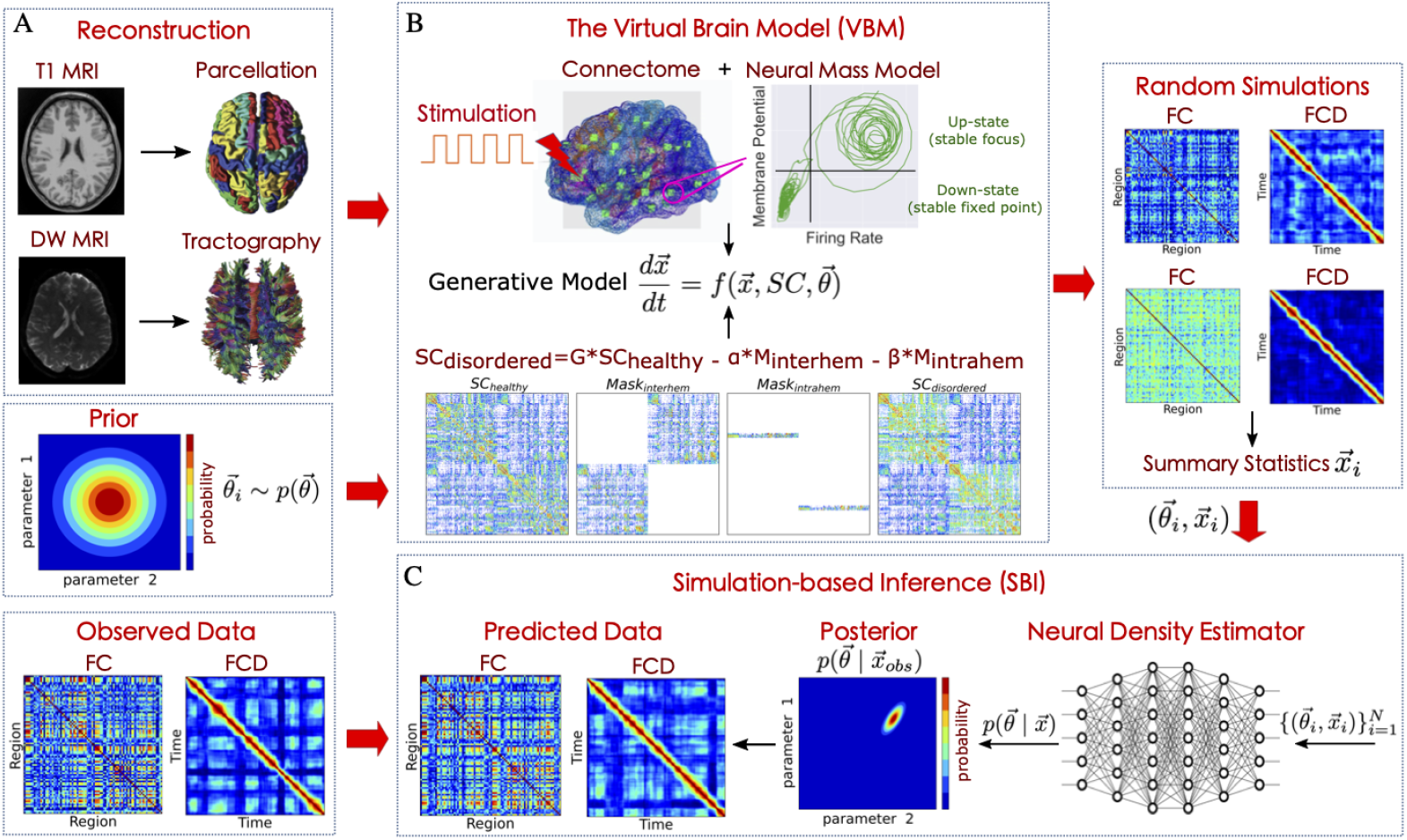
The SBI-VBMs workflow to estimate the posterior distribution of generative parameters in brain disorders. (**A**) TVB reconstruction pipeline. The T1-MRI and DW-MRI images are processed to build a personalized brain connectome. (**B**) The Virtual Brain Model (VBM). An exact neural mass model of spiking neurons is placed at each brain region, combined with anatomical data to generate various data features, such as FC and FCD. Estimating the generative parameters with perturbation in the SC using a masking approach, and stimulating certain regions, provides predictions on degradation in the connectome caused by disorders. (**C**) The SBI with deep neural density estimators. First, the model parameters are drawn randomly from a prior distribution. Then, the VBM simulator takes the parameters as input and generates summary statistics of the data as output. Next, a class of deep neural density estimators is trained on the pairs of random parameters and corresponding data features to learn the joint posterior distribution of the model parameters. Finally, we can quickly approximate the posterior for new data and make probabilistic predictions that are consistent with the observed data.

### 2.3 Functional connectivity dynamic

To quantify the communication between different brain regions and associated dynamics, we computed both static Functional Connectivity (FC) and Functional Connectivity Dynamics (FCD), which is the time-variant representation of FC and reflects the fluctuation of the covariance matrix over time (see Supplementary, Fig S1). From the simulated BOLD signals generated by TVB, we extracted FC matrix by calculating the Pearson correlation between the BOLD signals of any two brain regions. In order to track the time-dependent changes in the FC, we computed the windowed FCD, with length of sliding window τ = 30 *sec* and step size of *t*_*w+1*_ − *t*_*w*_ = 6 *sec* (Hansen et al., 2015; Arbabyazd et al., 2020):

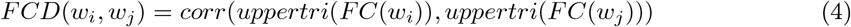

where, the (*i, j*) entry of the FCD matrix provided the Pearson correlation between the upper triangular parts of the two FC matrices *FC*(*w*_*i*_) and *FC*(*w*_*j*_) calculated at each sliding windows *w* = 1, 2, …, *N*_*w*_. To assess whether the simulated BOLD time-series might have state transitions between up and down states, we quantified the brain’s fluidity by calculating the FCD variance to capture the transitioning behavior (see Supplementary, Fig S1).

### 2.3 Bayesian inference

The Bayesian approach offers a principled way for making inference, prediction, and quantifying uncertainty for decision-making process (Gelman et al., 1995; Bishop, 2006; Box and Tiao, 2011). This approach naturally evaluates and incorporates uncertainties in the parameters and observations to drive meaningful conclusions from the data, with a broad range of applications (Bonomi et al., 2016; Hashemi et al., 2018; Khalvati et al., 2019; Guimerà et al., 2020; Broderick et al., 2023; Sip et al., 2023; Wang et al., 2023). Parameter estimation within a Bayesian framework is treated as the quantification and propagation of uncertainty through probability distributions placed on the parameters, updated with the information provided by data (Hashemi et al., 2020, 2021). Considering the joint probability distribution of the observation x and the unknown control parameters *θ*, referred to as the generative model. Then by the product rule: *p*(*θ, x*) = *p*(*θ | x*)*p*(*x*), equivalently, *p*(*θ, x*) = *p*(*x | θ*)*p*(*θ*). Hence, given data x and model parameters *θ*, Bayes rule defines the posterior distribution as

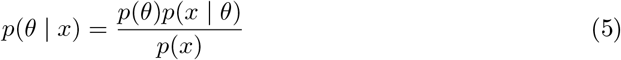

that combines and actualizes prior information (domain expertise) about unknown parameters with the knowledge acquired from observed data through the so-called likelihood function (data generating process). The prior information *p*(*θ*) is typically determined before seeing the data through beliefs and previous evidence about possible values of the parameters. The likelihood function *p*(*x | θ*) represents the probability of some observed outcomes given a certain set of parameters (the information about the parameters provided by the observed data). The denominator *p*(*x*) = ∫ *p*(*x | θ*)*p*(*θ*)*dθ* represents the probability of the data and it is known as evidence or marginal likelihood. In the context of inference, this term amounts to simply a normalization factor. Note that using Bayesian inference in this study, we aim to estimate the entire posterior distribution of the unknown parameters, i.e., the uncertainty over a range of plausible parameter values that generates the data, rather than a single point estimate for data fitting (e.g., the maximum likelihood estimation; Hashemi et al. (2018), or the maximum a posteriori; Razi et al. (2015); Friston et al. (2014); Vattikonda et al. (2021)).

### 2.5 Simulation-based inference

The key challenge to perform an efficient Bayesian inference is the evaluation of likelihood function *p*(*x | θ*). This is typically intractable for high-dimensional models involving non-linear latent variables (such as VBMs), as the likelihood evaluation requires an integral over all possible trajectories through the latent space that controls the generative process: *p*(*x | θ*) = *p*(*x, z | θ*)*dz*, where *p*(*x, z | θ*) is the joint probability density of data *x* and unmeasured (hidden) latent variables *z*, given parameters *θ*. In particular, when dealing with high-dimensional latent space at whole-brain scales, the computational cost of evaluating the likelihood function can become prohibitive. This makes likelihood-based approaches, such as Monte Carlo sampling, inapplicable (Hashemi et al., 2023). Simulation-Based Inference (SBI) or likelihood-free inference performs efficient Bayesian inference for complex models where the calculation of likelihood function is either analytically or computationally intractable (Cranmer et al., 2020). Instead of direct sampling from distributions and explicit evaluation of likelihood function, SBI sidesteps this problem by employing deep artificial neural networks (ANNs) to learn an invertible transformation between parameters and data (more precisely, between the features of a simulated dataset and parameters of a parameterized approximation of the posterior distribution).

Taking prior distribution 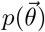 over the parameters of interest 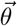, a limited number of *N* simulations are generated for training step 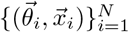, where 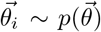 and 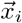 is the simulated data given model parameters 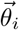. In other words, the training data set is an ensemble of *N* independent and identically distributed samples from the generative model 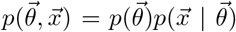, that can be run in parallel. After the training step, we are able to efficiently estimate the approximated posterior 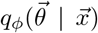 with learnable parameters *ϕ*, so that for the observed data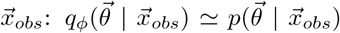. For more details, see Hashemi et al. (2023). Note that due to the amortized strategy, for any new observed data 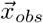, we can readily approximate the true posterior 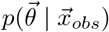 by a forward pass through the trained ANN (in the order of seconds).

Under preserving total probability used in this approach, the prior distribution (a simple distribution such as a uniform or standard normal) is transformed into the posterior distribution (a more complex potentially multi-modal), through a sequence of invertible transformation. This class of ANNs used for sampling and probability density estimation is called Normalizing Flows (NFs; Rezende and Mohamed (2015)). NFs are a family of generative machine learning models that convert a base distribution into any complex target distribution, where both sampling and density estimation can be efficient and exact (Papamakarios et al., 2019a; Kobyzev et al., 2020). The aim of density estimation is to offer an accurate description of the underlying probability distribution of an observable data set, where the density itself is unknown. In particular, NFs embedded in the SBI approach, such as (sequential) neural posterior estimation (Gonçalves et al. (2020); Lueckmann et al. (2017), allow for the direct estimation of joint posterior distributions, and bypass the need for MCMC, while also potentially capturing degeneracy or multi-modalities. Recently advanced NFs such as Masked Autoregressive Flows (MAF; Papamakarios et al. (2017)) and Neural Spline Flows (NSF; Durkan et al. (2019)) provide efficient and exact density evaluation and sampling from the joint distribution of high-dimensional random variables in a single neural-network pass. These generative models use deep neural networks to learn complex mappings between input data and their corresponding probability densities. They have achieved state-of-the-art performance with diverse applications, which efficiently represent rich structured and multi-modal posterior distributions, capturing complex dependencies and variations in the data distribution (Papamakarios et al., 2019a; Kobyzev et al., 2020). In this study, we use MAF and NSF models, which support invertible nonlinear transformations, and enables highly expressive transformations with low-cost computation. By training MAF/NSF on the virtual brain simulations with random parameters, we are able to readily estimate the full posterior of parameters from low-dimensional data features, such as FC/FCD.

### 2.6 Sensitivity analysis

Sensitivity analysis is a necessary step to determine which model parameters mostly contribute to variations in the model’s behavior due to changes in the model’s input, i.e., the identifiability analysis (Hashemi et al., 2018, 2023). A local sensitivity coefficient measures the influence of small changes in one model parameter on the model output, while the other parameters are held constant. This can be quantified by computing the curvature of objective function through the Hessian matrix (Bates and Watts, 1980; Hashemi et al., 2018):

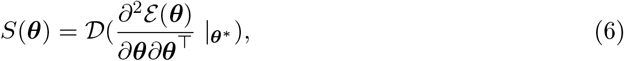

where *𝒟* denotes the main diagonal elements of a matrix, *ε* indicates a fitness function, centered at estimated parameters ***θ***^*∗*^.

Using the SBI approach, after the training step and posterior estimation for a specific observation, we can also efficiently perform sensitivity analysis by calculating the eigenvalues and corresponding eigenvectors from (Tejero-Cantero et al., 2020; Deistler et al., 2021):

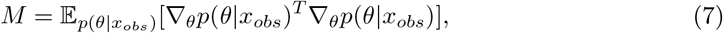

which then does an eigendecomposition *M* = *Q*Λ*Q*^*−1*^. A strong eigenvalue of the so-called active subspaces (Constantine, 2015) indicates that the gradient of the posterior is large, hence, the system output is sensitive to changes along the direction of the corresponding eigenvector.

### 2.7 Evaluation of posterior fit

To measure the reliability of the Bayesian inference using synthetic data, we evaluate the posterior z-scores (denoted by z) against the posterior shrinkage (denoted by s), which are defined as (Betancourt, 2014):

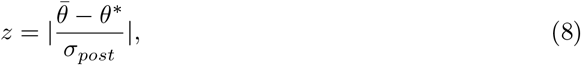

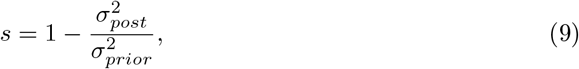

where 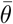 and *θ*^*∗*^ are the posterior mean and the true values, respectively, whereas *σ*_*prior*_, and σ_*post*_ indicate the standard deviations of the prior and the posterior, respectively. The posterior z-score quantifies how much the posterior distribution of a parameter encompasses its true value. The posterior shrinkage quantifies how much the posterior distribution contracts from the initial prior distribution. The concentration of estimations towards large shrinkages indicates that all the posteriors are well-identified, while the concentration towards small z-scores indicates that the true values are accurately encompassed in the posteriors.

## 3. Results

### 3.1 The SBI-VBMs workflow

Figure 1 illustrates the overview of our approach, referred to as Simulation-Based Inference on Virtual Brain Models (SBI-VBMs), to make probabilistic predictions on brain disorders by estimating the joint posterior distribution of the generative parameters. This approach relies only on random model simulations to efficiently approximate the posterior distribution of unknown parameters, without requiring exact likelihood evaluation.

At the first step, the non-invasive brain imaging data such as T1-weighted MRI and Diffusion-weighted MRI (DW-MRI) are collected for a specific patient (Fig 1**A**). The T1-weighted MRI images are processed to obtain brain parcellation, and diffusion-weighted (DW-MRI) images are used for tractography. With the generated fiber tracts and the regions defined by the brain parcellation, the connectome (i.e., the total set of links between brain regions) is constructed by counting the fibers connecting all regions. The SC matrix, whose entries represent the connection strength between brain regions, provides the basis for constructing a network model–the virtual brain– which is capable of generating various functional brain imaging data at arbitrary locations (e.g., cortical and subcortical structures). This step constitutes the structural brain network component, which imposes a constraint on brain network dynamics, i.e., the evolution of trajectories in the latent space, allowing the hidden state dynamics to be inferred from the data.

Then, each brain network node is assigned a computational model of average neuronal activity, a so-called neural mass model (see Fig 1**B**). Here, we use an exact mean-field approximation for a population of spiking neurons (see Eq. (1)), which due to its rich dynamics, such as bi-stability, enables us to generate various spatio-temporal dynamics as observed experimentally. This combination of the brain’s anatomical information with the mathematical modeling of averaged dynamics at the level of local neural populations (e.g., 16 cm^2^ of the cortical surface) constitutes the functional brain network component, characterizing the communication between brain regions.

Subsequently, the perturbation of SC by spatial masking, the inter- and intra-hemispheric edges can provide an estimation of the level of degradation in the patient connectome caused by disorders (see Eq. (3)). Notably, the stimulation of certain brain regions can be used to increase the causal evidence in the structure-function relationship. Embedding the disordered SC into the VBM, then complements the generative brain model of disorders (see Eq. (2), as a nonlinear dynamical model 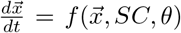, with control parameters *θ*). The set of generative parameters comprises the regional excitability parameters *η*_*i*_, a global coupling parameter *G* describing the scaling of the brain’s *SC*, and parameters *α* and *β* indicating the levels of interhemispheric deterioration, and intrahemispheric degradation within the limbic system, respectively. With an optimal set of these parameters, VBMs can mimic essential data features observed in fMRI recordings, such as FC/FCD matrices.

Finally, we use SBI (see Fig 1**C**) to estimate the posterior distribution of the generative parameters in brain disorders. To perform SBI with deep neural density estimators, we first draw the parameters randomly from a prior distribution, i.e., 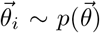, for each *i*-th simulation. The VBM simulator takes the parameters 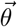 as input and generates summary statistics 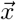 of the data as output. By preparing a training dataset through performing VBM simulations with random parameters, a class of deep neural density estimators (such as MAF or NSF model) is then trained on 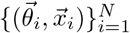 with budget of N simulations, to learn an invertible transformation between data features and parameters of an approximated posterior distribution 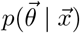 (see Eq. (5)).

Notably, for any new observation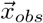, we can readily approximate the posterior 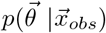, making predictions that are probabilistically consistent with the observed data. This approach allows for efficient and accurate inference of the full posterior, with no demand for further simulations at the inference step due to the amortization strategy (i.e., a single pass through the ANNs). This is the primary advantage of this approach that the amortized inference at the subject level enables us to evaluate different hypotheses with uncertainty quantification for decision-making processes at a negligible computational cost, typically in the order of a few seconds.

### 3.2 Sensitivity analysis on the degree of brain degradation

First, we perform a sensitivity analysis on the degree of perturbation in the connectome (see Eq. (2) and Eq. (3)). The sensitivity analysis can shed light on the identifiability of model parameters. A small change in a very sensitive model parameter causes a strong response in the model output, which indicates that the parameter is more identifiable. On the contrary, a model parameter with low sensitivity is more elusive to identify, because any modification in an insensitive parameter has no influence on the model output. To assess the sensitivity of the parameters, we used the Hessian matrix (see Eq. (6)), a metric describing the local curvature of a function based on its second partial derivatives.

Figure 2**A**-**C** shows the Kolmogorov-Smirnov (KS) distance between the distributions of values in the observed and predicted FC matrices (in blue) and FCD matrices (in red), for a sweep over the global coupling parameter *G*, and the levels of inter- and intra-hemispheric degradation denoted by *α* and *β*, respectively. The results indicate a unique global minimum in the objective function defined by the KS distance, but with varying sensitivity values as indicated by the Hessian matrix in Fig 2**D**. Note that the finite confidence intervals in the profile of the *β* mask indicate a practical non-identifiability rather than structural non-identifiability (which demonstrates a flat valley that extends infinitely in both the upper and lower bounds).

**Fig. 2.**
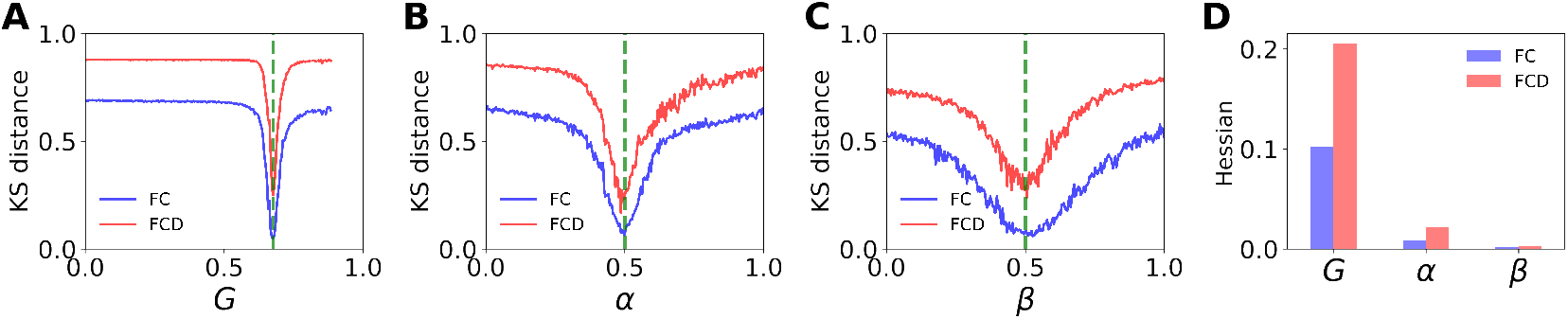
Sensitivity analysis on the degree of degradation in a personalized connectome caused by virtual disorders. The plots represent the results of a grid search over the KS distance between the distribution of observed and predicted FC (in blue) and FCD (in red), computed for incremental increase in (**A**) the global scaling parameter denoted by *G*, (**B**) the degree of degradation in inter-hemispheric connections denoted by *α*, and (**C**) the level of intra-hemispheric degradation within the limbic system, represented by *β*. (**D**) The Hessian values, quantifying the local curvature of KS distance at the ground truth values (in green), indicate more sensitivity to FCD than FC. The sensitivity of the *β* mask is nearly zero (see Tabel S1).

Interestingly, the generative parameters show greater sensitivity to FCD compared to FC (see Tabel S1), while the sensitivity to *β* mask, representing the level of intra-hemispheric degradation (within the limbic system), is near zero, indicating non-identifiability for its estimation. See Supplementary Fig S3 for similar results using Kullback-Leibler (KL) divergence between the distributions of observed and predicted FC and FCD matrices. Supplementary Fig S4 shows the KL divergence between these features at the optimal points. Note that reporting an error metric between predicted and observed BOLD time-series, such as the root mean square error, can be misleading due to the high noise present in the generated signals.

An important question that arises is how to effectively integrate spatio-temporal and functional data features into the inference process. This involves incorporating relevant information derived from both FC and FCD, including summary statistics such as variance (fluidity) as well as the (high-order) statistical moments of BOLD data, beyond the KS distance (synchronization). In addition, we need to assess the uncertainty associated with the estimated parameters and their relationship to ascertain parameter identifiability. These questions will be addressed by SBI, which enables us to naturally account for uncertainty and identifiability concerns while leveraging the maximum information available from the data for accurate and reliable inference.

### 3.3 Inference on level of degradation in connectome

Here we use SBI to estimate posterior distribution in the set of unknown generative parameters denoted by 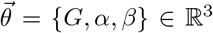. Figure 3 shows the estimated posterior using SBI, by investigating the impact of functional data features (FC and/or FCD), as well as the number of simulations in quantifying posterior uncertainty. For the training (here using MAF model), the parameters are drawn from a uniform prior in the range: *θ*_*i*_ ∈ 𝒰 (0, 1). It can be seen that the SBI accurately estimates the posterior of parameters G and α using functional data features from a budget of 100k simulations (see Fig 3**A, B**). However, we observed no posterior shrinkage in the estimation of *β* mask (see Fig 3**C**). This is due to the model’s insensitivity to intra-hemispheric deterioration compared to the whole-network and intra-hemispheric degradation (see Fig 3**D**), as calculated from the eigenvalues of the posterior density (see Eq. 7). This is in agreement with the sensitivity analysis conducted using the Hessian matrix (see Fig 2**D**).

**Fig. 3.**
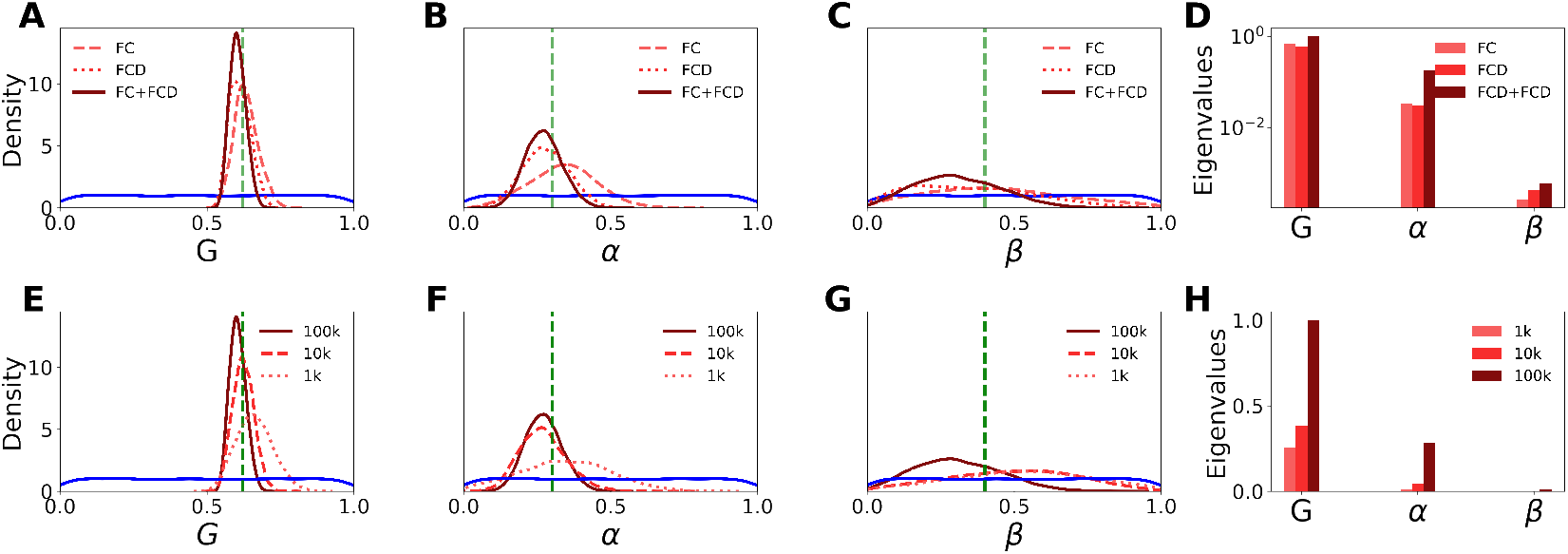
Inference on the global, inter- and intra-hemispheric degradation in a virtual brain. The posterior distribution estimated using 100k random simulations for training on functional data features (FC and/or FCD), is shown (in red) for (**A**) the global coupling parameter *G*, (**B**) the inter-hemispheric deterioration *α*, and (**C**) the intra-hemispheric degradation *β*. The parameters are drawn from a uniform prior between zero and one (in blue). The ground truth values are *G* = 0.62, *α* = 0.3, *β* = 0.4 (in green). (**D**) The sensitivity analysis conducted using active subspaces explains the lack of posterior shrinkages in the estimation of *β*. (**E**), (**F**), (**G**) The estimated posteriors of parameters *G, α*, and *β*, respectively, by increasing thee simulation budget for training. (**H**) As the number of simulations for training increases, the sensitivity increases for parameters *G*, and *α*, but not for *β*, indicating the non-identifiability of degradation within the limbic system.

Moreover, the uncertainty estimation indicates that FCD is more informative than FC when inferring perturbed connections. However, using both data features provides even stronger evidence against the fitted data (see solid lines in Fig 3**A**-**D**). We also observe that increasing the number of simulation budget during the training step provides more informative posterior distributions for parameters *G* and *α*, but not for parameter *β* (see Fig 3**E**-**H**). These results demonstrate the non-identifiability in estimating the intra-hemispheric degradation from functional data features, even when a large number of model simulations are used for training.

The effect of the simulation budget on the uncertainty of posterior when training with the state-of-the-art deep neural density estimators (MAF and NSF) is shown in Fig S5. We observed that both MAF and NSF models are compatible in uncertainty quantification (with different simulation budgets), but MAF was 2-4 times faster than NSF during the training process.

### 3.4 Inference on perturbation-based degradation in connectome

The structural non-identifiability is independent from the accuracy of experimental data. Simply increasing the quantity or quality of existing measurements will not resolve this issue.

The remedy is to design a new setup for the measurements (e.g., new mapping function from the source to sensor space or reparameterizing the model configuration space). In contrast, the practical non-identifiability arises due to the limited amount and/or quality of observations. The remedy for this issue is to add more data so that it provides sufficient constraining power in the latent space dynamics, yielding a unique estimation with finite confidence intervals.

To address the practical non-identifiability issue in estimating the *β* mask, we included the statistical characteristics of the BOLD time-series (such as the mean, variance, skewness, and kurtosis of BOLD time-series) in the training step. As shown in Fig S6, adding such spatio-temporal data features results in the better estimation of model parameters, particularly in the estimation of *β* mask. However, this leads to a high correlation between the global scaling parameter *G* and the *α* mask (*ρ* = 0.9, see Fig S6**B**). Furthermore, the spatio-temporal data features such as mean and variance can become redundant when using signal processing techniques such as normalization or z-scoring. To address this issue, we have perturbed the brain dynamics through stimulation. Fig 4 illustrates the consequence of brain stimulation on the recovery of generative parameters denoted by 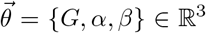. Here we used only functional data features for training in SBI.

**Fig. 4.**
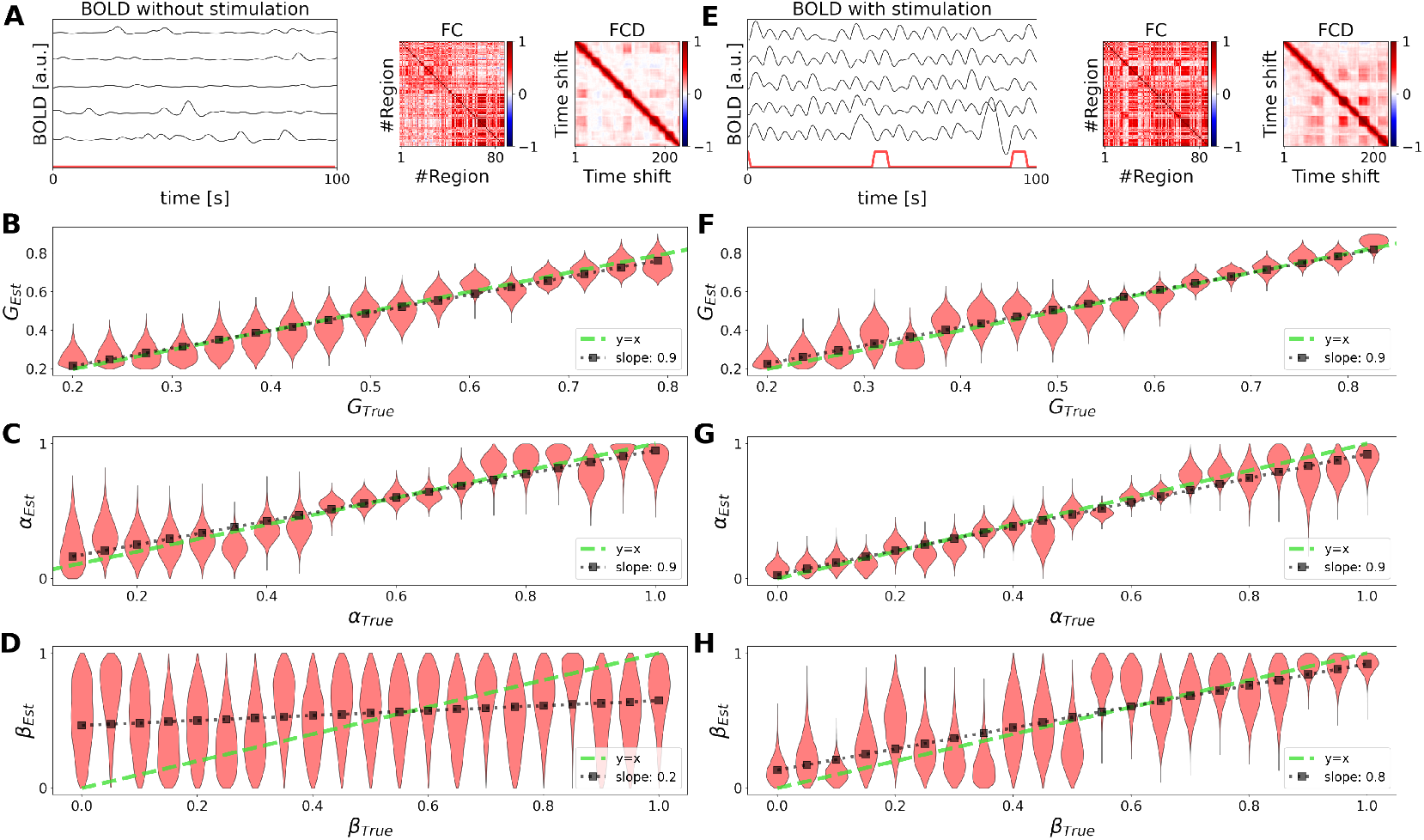
Inference on the global, inter- and intra-hemispheric degradation in a virtual brain with perturbed dynamics induced by stimulation. Here we used only functional data features for training in SBI. (**A**), (**E**) The simulated BOLD data and corresponding FC and FCD matrices, under conditions of no-stimulation and with-stimulation, respectively. (**B**), (**F**) SBI provides accurate recovery of the parameter *G*, pre- and post-stimulation, respectively. (**C**), (**G**) Accurate posterior estimation of the *α* mask, before and after stimulation, respectively. (**D**), (**H**) SBI on *β* mask shows poor estimation when the stimulation is off, but reliable estimation when the stimulation is on, respectively. The violin plots in red show the estimated posterior densities. The black dots represent a linear regression on the maximum values of the estimated posteriors. Dashed green lines represent a perfect fit (ground truth values).

Fig 4**A, E** show the simulated BOLD data and the corresponding FC and FCD matrices, in the absence and presence of intervention by stimulation, respectively. It can be seen that the BOLD data display enhanced structural patterns, as evidenced by the higher FC and the increased variance in FCD (the fluidity before stimulation was 0.001, and it increased to 0.004 after stimulation). Here the stimulation is induced by adding a repetitive step current, characterized by a duration of 5 *sec* and an amplitude of 1.2 *μA*/*cm*^2^ to the membrane potential variable (through *I*_*stim*_ in Eq. (2)), within the limbic system.

Fig 4**B, F** show that SBI provides accurate recovery of global scaling factor *G*, before and after stimulation, respectively. This is consistent with our previous results indicating the model’s heightened sensitivity to this parameter. Fig 4**C, G** demonstrate the robustness of estimation on the *α* mask by closely matching the true values, regardless of the stimulation. This indicates the reliability of SBI in estimating the level of degradation between hemispheres, even by perturbation in brain dynamics.

Fig 4**D, H** show that SBI provides poor estimation on the *β* mask when the stimulation is off, but reliable estimation when the stimulation is applied, respectively. This demonstrates that the intervention by stimulation is able to address the non-identifiability issue encountered when estimating degradation within brain hemispheres. In sum, perturbing brain dynamics through stimulation increases the likelihood of transitions between brain states, thereby improving the quality of estimation by providing new observable responses for the inference process. Nevertheless, it may be impractical to administer stimulation to all subjects.

In the following, we extend the dimensionality of generative parameters to include the excitability parameter. We investigate different forms of model parameterization, such as pooled (homogeneous), unpooled (heterogeneous), and hierarchical (multi-level), to improve the efficiency of SBI-VBMs in higher dimensions, with no need for stimulation.

### 3.5 Inference on homogeneous generative parameters

Pooled modeling is a statistical technique that assumes a common structure or distribution across all individual units, treating them collectively as a single unit and assuming no variation in the sampling process. For simplicity purposes, homogeneous (or pooled) modeling of excitability assumes a uniform and shared excitability parameter across all brain regions.

Here we use SBI to estimate posterior distribution in the set of unknown generative parameters denoted by 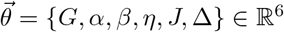. In this case, we infer the brain dynamics from parameters located in multiple brain regions with similar or homogeneous characteristics (see Supplementary, Fig S7**A**). This form of model parameterization reduces the spatial dimensions, hence the computational complexity of the inference process.

Fig 5 shows the estimated posterior distributions using both spatio-temporal and functional data features extracted from BOLD time-series. The diagonal panels show that the ground-truth values (in green vertical lines) used for generating observed data are well under the support of the estimated posterior distributions (in red). Here, we used 100k random simulations from uniform priors (in blue) as: *G, α, β ∈ 𝒰* (0, 1), *η ∈ 𝒰* (*−*6, *−*3.5), J *∈ 𝒰* (1, 30), and Δ *∈ 𝒰* (0.1, 2). Due to sufficient budget of simulations and the informativeness of the data features for training, hence the accurate parameter estimation, we can see a close agreement between the data features in the observed and predicted BOLD data (see FC/FCD matrices, shown in the lower diagonal panels). The joint posterior between parameters are shown at the upper diagonal panels, along with their correlation values (*ρ* shown at the upper left corners). It can be seen that there is a strong correlation between parameters *η* and *J* (with *ρ*=-0.9), also between *η* and Δ (with *ρ*=-0.9). This indicates a parameter degeneracy between the excitability and the spread of their distribution, as well the synaptic weight at whole-brain level. This can be due to the homogeneous parameterization used for inference, which ignores all variation among the units being sampled. Using MAF model, the training took around 45 *min*, whereas generating 10000 samples from the posterior took less than 2 *sec*. Supplementary Figure S8 demonstrates that excluding the spatio-temporal data features (i.e., statistical characteristics of BOLD time-series) during training leads to poor parameter estimation.

**Fig. 5.**
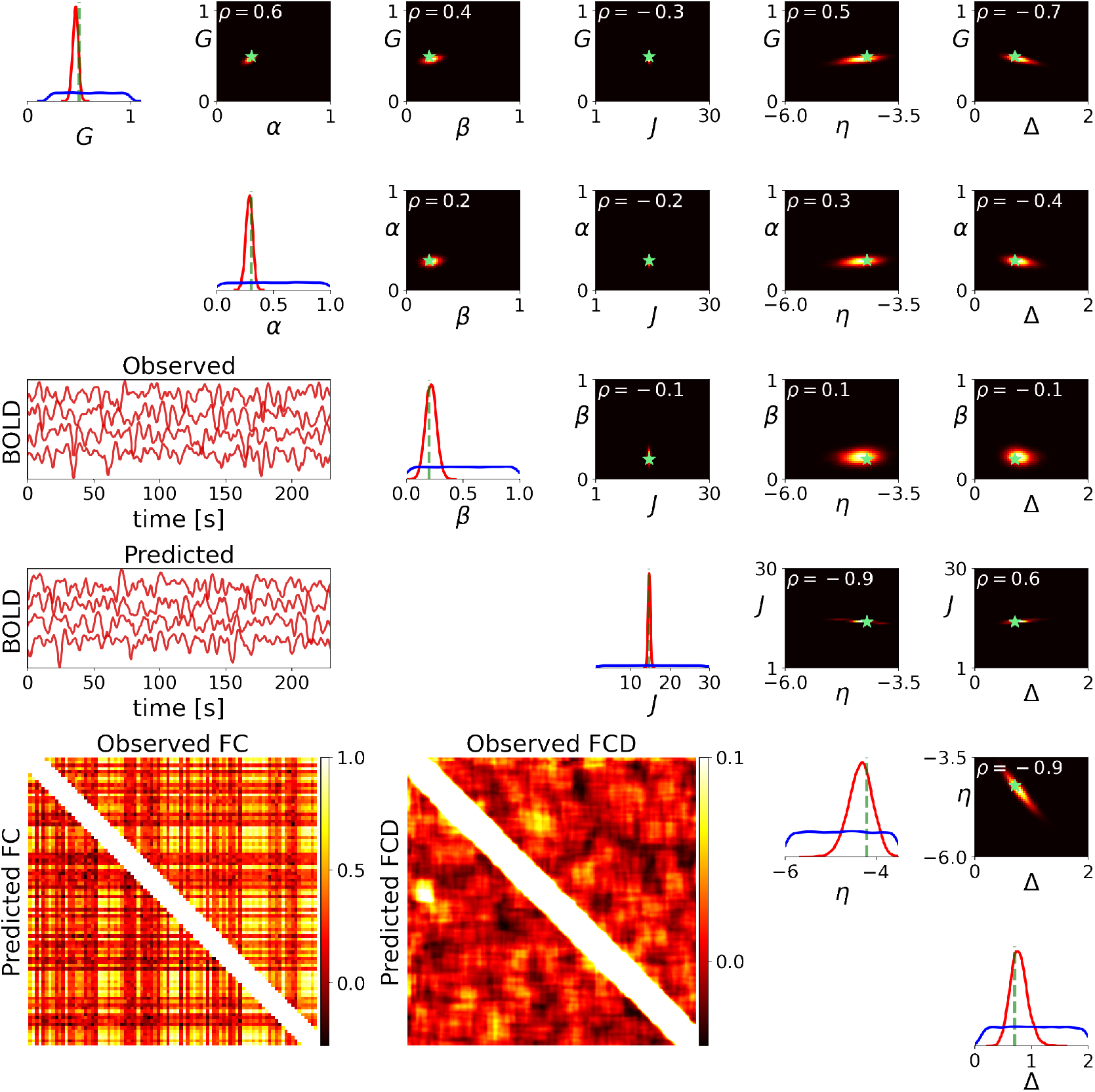
Inference on homogeneous generative parameters in a virtual brain. Here the set of inferred parameters is 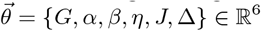. The training process involved using both spatio-temporal and functional data features, with a budget of 100k simulations. The diagonal panels display the prior (in blue) and estimated posterior (in red). The true values (green lines) are well under the support of the estimated posterior distributions. The lower diagonal panels display the observed and predicted BOLD data and the corresponding FC/FCD matrices. The upper diagonal panels show the joint posterior between parameters, and their correlation values (*ρ* at the upper left corners). The ground-truth values are shown by green stars. High-probability areas are color-coded in yellow, while low-probability areas are represented in black.

### 3.6 Inference on heterogeneous generative parameters

Unpooled modeling is a statistical technique that treats individual units as being sampled independently, allowing for adaptation to diverse characteristics and behaviors within the dataset, without assuming a common structure. Heterogeneous (or unpooled) modeling of excitability captures the distinct characteristics of each region without imposing a uniform structure at whole-brain level. This approach leads to more precise and biologically plausible inference on the brain’s (dis)functioning, but significantly increases the dimension of the parameter space.

Here we use SBI to estimate posterior distribution in the set of unknown generative parameters denoted by 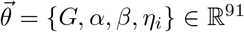, where *η*_*i*_ with *i ∈ {*1, 2, …, *N*_*n*_ = 88*}* is the excitability parameter heterogeneously distributed across brain regions (see Supplementary, Fig S7**B**). This form of model parameterization offers a comprehensive explanation for the variation in the excitability across brain regions. Fig 6 shows the maximum of estimated posterior distribution for three different scenarios in setting the excitabilities across brain regions. It can be seen that SBI provides accurate estimation for different configurations in excitabilities across brain regions. Here, the training process involved using both spatiotemporal and functional data features. See Supplementary, Fig S9 for the setups and full estimated posteriors. Importantly, using only the functional features (FC and FCD) does not provide sufficient information to infer heterogeneous excitability across whole-brain regions (see Fig S10).

**Fig. 6.**
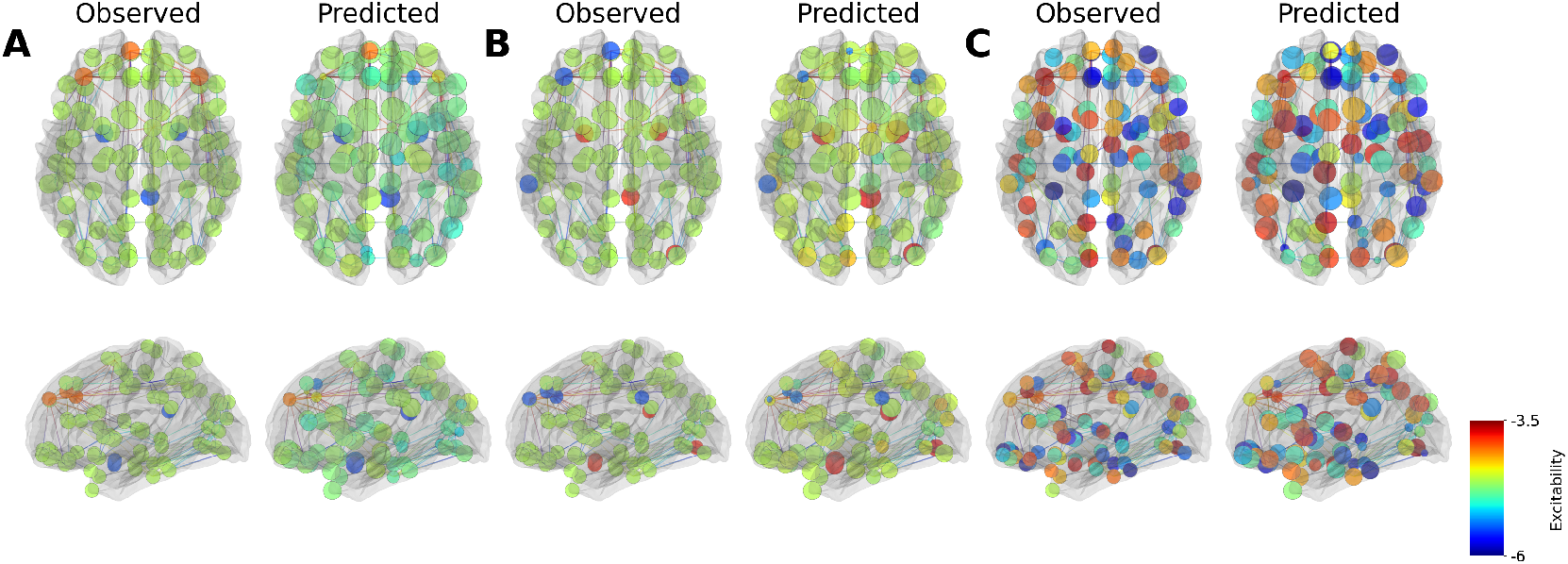
Inference on heterogeneous generative parameters in a virtual brain. Here the set of inferred parameters is 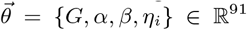. SBI provides accurate posterior estimation for various scenarios by setting different excitability maps across brain regions. Both spatio-temporal and functional data features were used for training, with a budget of 1M simulations. The regions are color-coded based on their excitability values, with blue representing low-, red representing medium-, and green representing high-excitability. At top: axial view. At bottom: sagittal view.

Regarding the computational cost, using MAF model, the training on 1M simulations took around 10 *h*, whereas generating 10000 samples from the posterior took less than 60 *sec*. Figures S11, S12 demonstrate that both MAF and NSF models were compatible in uncertainty quantification of heterogeneous parameters. Note that here we used 1M random simulations, compared to the 100k simulations used in homogeneous parameterization (Fig 5). This indicates the demand for a very large simulation budget during training step to achieve accurate estimation based on a heterogeneous parameterization.

### 3.7 Inference on hierarchical generative parameters

Hierarchical (or partial pooling) models offer a middle ground between the simplicity of pooled models (which assume homogeneity) and the complexity of fully unpooled models (which assume complete independence by heterogeneity). This balance allows for more flexibility in modeling, and more efficiency and accuracy in estimating.

Here we use SBI to estimate posterior distribution in the set of unknown generative parameters denoted by 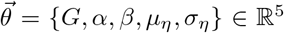, where 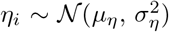, indicating that excitabilities are sampled from a Gaussian distribution with mean centered at *μ*_*η*_, and standard deviation of *σ*_*η*_. This is a hierarchical parameterization on excitabilities (see Supplementary, Fig S7**C**), in which they are sampled from a population distribution so that the heterogeneous parameters *η*_*i*_ with *i ∈ {*1, 2, …, *N*_*n*_} are now sampled from a generative distribution *𝒩* (*μ*_*η*_, *σ*_*η*_). Hence, only two hyperparameters μ_η_ and σ_η_ need to be estimated to explain the variation in excitability across the brain, rather than N_n_ excitability parameters.

Fig 7 shows the estimated posterior by SBI using the both spatio-temporal and functional data features for training. The diagonal panels show that the posterior (in red) accurately encompass the ground-truth values (in green vertical lines) used for generating the observed data. For training, we generated 100k random simulations from uniform priors (in blue) as: *G, α, β ∈ 𝒰* (0, 1), *μ*_*η*_ ∈ *𝒰* (*−*6, *−*4), and *σ*_η_ *∈ 𝒰* (0, 1). Due to effective form of model parameterization, hence the accurate parameter estimation, we can see a close agreement between the data features in the observed and predicted BOLD data, such as the FC/FCD matrices (shown at the lower diagonal panels). The joint posterior between parameters are shown at the upper diagonal panels, along with their correlation values (*ρ* at the upper left corners). Using MAF model, the training took around 30 *min*, whereas generating 10000 samples from the posterior took less than 1 *sec*. In contrast to heterogeneous modeling, the hierarchical structure yields more informative posteriors when using only functional data features, even with a significantly smaller budget of simulations for training (see Supplementary, Figure S10 versus Figure S13). These results indicate that hierarchical approach effectively takes into account the variability across brain regions while naturally incorporating a form of regularization to provide more stable and robust estimates.

**Fig. 7.**
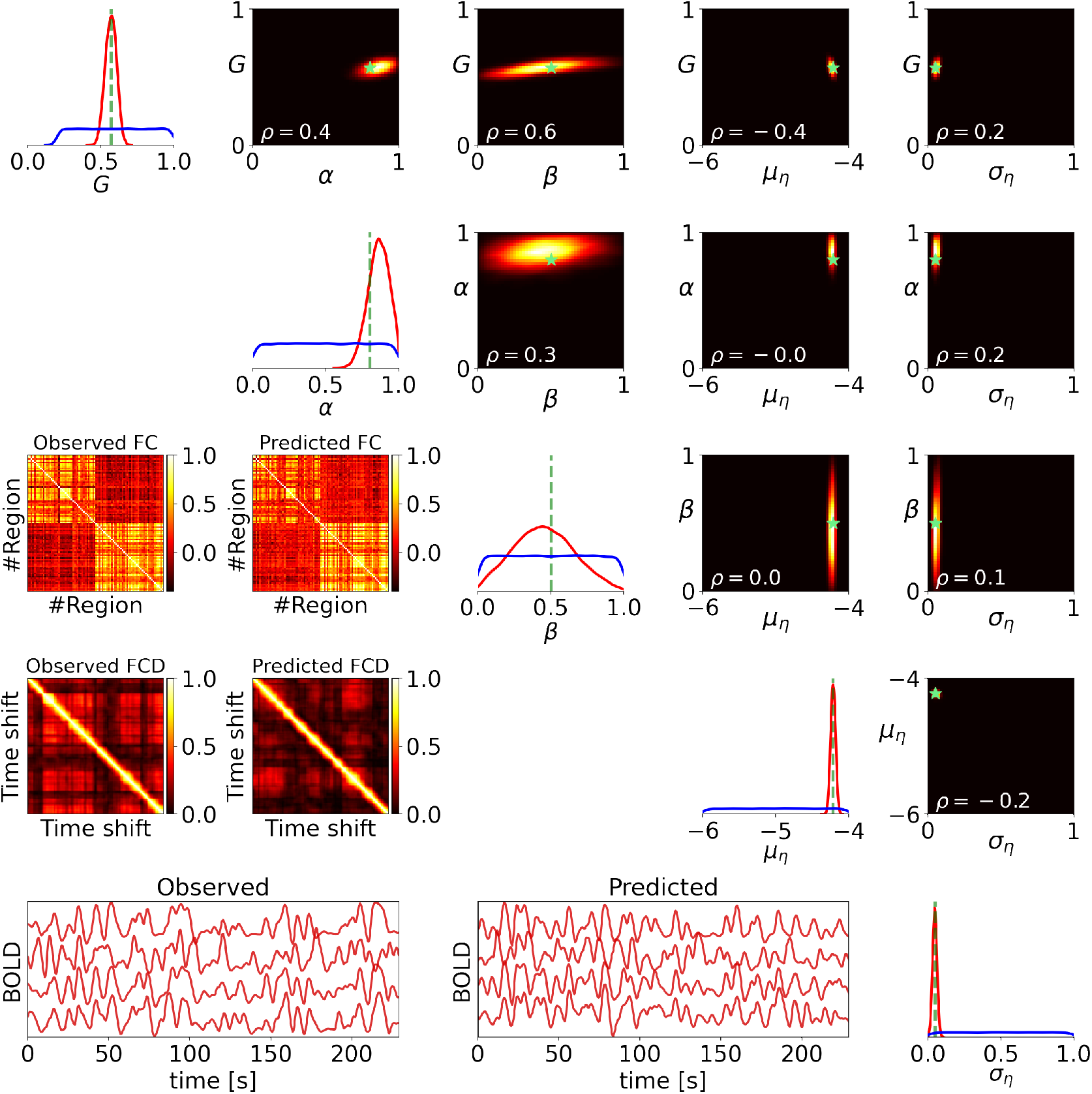
Inference on hierarchical generative parameters in a virtual brain. Here the set of inferred parameters is 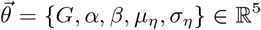, with 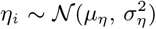. We used both spatio-temporal and functional data features, for the training on a budget of 100k simulations. The diagonal panels display the prior (in blue) and estimated posterior distributions (in red). The true values (green lines) are well under the support of the estimated posterior distributions. The lower diagonal panels display the observed and predicted BOLD data and the corresponding FC/FCD matrices. The upper diagonal panels show the joint posterior between parameters, and their correlation values (*ρ* at the lower left corners). The ground-truth values are shown by green stars. High-probability areas are color-coded in yellow, while low-probability areas are represented in black.

## 4. Discussion

This study addresses the ongoing challenge of probabilistic inference on complex wholebrain dynamics using connectome-based models, by presenting simulation-based inference on virtual brain models (SBI-VBMs). Our methodology (see Fig 1) offers a compelling advantage in terms of parallel and fast simulations, especially when utilized with the computational power of GPUs (see Supplementary, Fig S2). We showed that by systematically examining how variations in input parameters affect the output, sensitivity analysis (see Fig 2, and Fig S3) is crucial in ensuring the robustness and reliability of inference results, thus for decision-making processes. Our results underscore the limitations of relying simply on functional data features for making inferences about degradation in brain connectivity and regional excitability (see Figs 3-4, and Supplementary, Fig S10). Rather, we proposed the SBI-VBMs approach, which relies on expressive deep neural networks to easily incorporate all relevant information, such as spatio-temporal and functional features. This approach demonstrates its effectiveness in accurately estimating generative parameters associated with brain disorders (see Figs 5-7). Furthermore, perturbing the brain dynamics through stimulation, which increases the likelihood of transitions between brain states, provides a valuable tool for resolving the issue of non-identifiability, especially in the context of intra-hemispheric connections (see Fig 4). Ultimately, the hierarchical structure of SBI-VBMs emerges as an efficient parameterization, enabling precise and biologically meaningful inference on brain’s (dys)functioning (see Fig 7, and Fig S13).

Several previous studies in whole-brain modeling (Deco et al., 2014; Wang et al., 2019; Kong et al., 2021; Cabral et al., 2022) have used optimization methods to provide a single best value of an objective (or a cost) function, scoring the model performance against the observed data (such as minimizing Kolmogorov-Smirnov distance or maximizing the Pearson correlation between observed and generated FC/FCD). Such a parametric approach results in only a point estimation, and is limited to construct the relationship between parameters and the associated uncertainty (see Fig 5, and Fig 7). The optimization algorithms may easily get stuck in local extrema, requiring multi-start strategies to address the potential multimodalities and parameter degeneracies. Moreover, the estimation depends critically on the form of objective function defined for optimization (Svensson et al., 2012; Hashemi et al., 2018), and the models involving differential equations often have strongly correlated and/or non-identifiable parameters (Raue et al., 2009; Wieland et al., 2021). These issues can be addressed by using Bayesian inference which provides proper representation of distributions and dependence among parameters (Samaniego, 2010; Hashemi et al., 2018).

Bayesian approach has the advantage of yielding a joint probability distribution for model parameters (see Supplementary, Fig S6, and Figs S8-S13). The posterior distribution encompasses all possible parameter combinations that produce a simulation output that explains the data with quantifying the uncertainty in estimation, which is critical for model comparison, selection and averaging, hypothesis testing, and decision-making process. Bayesian inference also allows us to integrate the prior knowledge in inference and prediction (Gelman et al., 2014) e.g., the physiological, anatomical, or clinical knowledge to maximize the model predictive power against measurements (Hashemi et al., 2021; Baldy et al., 2023).

Learning the parameters of dynamical systems enables one to infer not only beliefs or probabilities of events but also the dynamics of events under changing conditions, for instance, changes induced by treatments or external interventions (Friston et al., 2003; Pearl, 2009). A well-established Bayesian framework for inferring hidden neural state dynamics from neuroimaging data is the so-called Dynamical Causal Modeling (DCM; Friston et al. (2003)). Our SBI approach shares a key aspect with DCM: Bayesian inference for causal effects by estimating the posterior distribution of parameters of a dynamical generative model. However, there are key differences in practice. The connectome-based approach used in this study considers the structural connectivity as fixed parameters which are obtained from non-invasive imaging data of individuals. In contrast, DCM relies on effective connectivity to explain the effects on observation induced by the causal changes in interactions among brain regions (Frässle et al., 2017, 2018). Although effective connectivity can provide a better model fit to empirical data due to the high-dimensionality of parameter space, it may easily suffer from the non-identifiability issue, unless the changes in connectivity are constrained by the prior belief that there are transitions among a small number of brain connectivity states (Zarghami and Friston, 2020). In DCM, the non-linear ordinary differential equation representing the neural mass model is often approximated by its linearization around the system fixed points, whereas the non-linear property of generative models to ensure a switching behavior in the data is maintained in our approach (see Eq. (2)). More importantly, inversion of a DCM involves minimizing the free energy (equivalently, the Kullback-Leibler divergence), in order to maximize the model evidence (Friston et al., 2014; Razi et al., 2015), while its mean-field variant ignores - by definition-the correlation between parameters. Our approach is equipped with state-of-the-art deep learning algorithms for Bayesian inference (e.g., MAF/NSF models), in which systematically place a tighter upper bound on model evidence (Rezende and Mohamed, 2015; Papadopoulou et al., 2017) to deal with potential muti-modalities and degeneracies among parameters (Hashemi et al., 2023).

Recently, we have proposed a whole-brain probabilistic framework, the so-called Bayesian Virtual Epileptic Patient (BVEP; Hashemi et al. (2020)), in which the generative model based on a system of high-dimensional and nonlinear stochastic ordinary differential equations was inverted using unbiased and automatic MCMC sampling algorithms. Although, we have shown that this non-parametric approach is able to accurately estimate the spatial map of epileptogenicity across whole-brain areas, it required a reparameterization over model configuration space to facilitate efficient exploration of the posterior distribution in terms of computational time and convergence diagnostics (Jha et al., 2022). In the presence of metastability in the state space (see Fig 1, Fig S1), MCMC methods require either more computational cost or intricately designed sampling strategies (Gabrié et al., 2022; Jha et al., 2022; Baldy et al., 2023), whereas SBI allows for efficient Bayesian estimation without the access to the full knowledge on the state space representation of a system (Baldy et al., 2024). Note that we used the data features derived from only firing rates, while the information related to membrane potential activities was treated as missing data (i.e., no access to the full state space behavior). More critically, this is highly beneficial if the model output such as simulated raw time-series poses discrepancy with the observed data, rather, it accurately explains the low-dimensional data features such as FC/FCD. Providing fast simulations, SBI can be applied to other whole-brain network models, since it requires neither model nor data features to be differentiable (Gonçalves et al., 2020; Hashemi et al., 2023). Nevertheless, finding low-dimensional but sufficient informative that can deal with parameter degeneracies, the noise estimation, and scalability to higher dimension with a feasible budget of simulations are the challenges for this approach to be applied on other neuroimaging datasets.

In recent work by Sip et al. (2023), a data-driven approach has been introduced to infer both the unknown generative dynamical system and the parameters varying across brain regions, while network nodes representing the brain regions are connected with realistic strengths derived from diffusion-weighted imaging data. This method offers a different perspective, leveraging the power of deep generative moodels (Variational Autoencoders) to focus on the role of regional variance of model parameters. This approach bypasses the need for specific mathematical form of the neural mass models at each region, allowing for increased flexibility and the potential to capture complex relationships within neuroimaging data. In contrast, the current study employs an exact mean-field description of spiking neurons to ensure both realistic simulations of brain activities and the interpretability of the estimations to develop effective interventions. However, determining causality in brain disorders requires careful consideration of multiple variables, including longitudinal data, multimodal imaging techniques, and consistency of statistical and computational methods with the data. Moreover, brain disorders often involve intricate interactions between genetic, environmental, and neurobiological factors, making it challenging to isolate specific causal factors.

In summary, the methodology employed in this study offers the advantage of parallel and fast simulations for flexible, scalable, and efficient probabilistic inference and can be applied to different connectome-based models. The key challenge lies in identifying low-dimensional yet informative data features that can effectively deal with the non-identifiability issues in the generative models. Nevertheless, hierarchical structures serve as an effective solution, especially in cases where informative data features or perturbing brain dynamics are not easily available. The applications of our approach to inferring the origin of brain diseases remain to be explored in future studies. This investigation holds promise for advancing our understanding and potentially informing targeted interventions to clinicians for different brain disorders.

## 5. Technical Terms

### Bayesian Rule

A fundamental belief updating principle that calculates the probability of a hypothesis given new evidence.

### Connectome

The total set of links between brain regions.

### Deep neural density estimators

A class of neural network-based approaches that are used to learn and approximate the underlying probability distribution from a given dataset.

### Evidence

Empirical data or observable information that can be used to support or reject a hypothesis.

### Generative parameters

The setting or configuration within a generative model that controls the synthesis of data and represents causal relationships.

### Generative model

A statistical, machine learning, or mechanistic model that represents the underlying data distribution to generate new data resembling the original dataset.

### Hierarchical modeling

A statistical modeling approach where multiple levels of probability distributions are used to represent uncertainty and variability in a complex system, with structured dependencies.

### Hidden state

A possible model outcome describing system behaviour, which cannot be measured directly but only inferred indirectly through observed data.

### Hypothesis

A statement on potential causality or predictive relationships to observed data.

### Likelihood

The conditional probability of observing the evidence given a particular hypothesis.

### Markov chain Monte Carlo (MCMC)

A family of stochastic algorithms used for uncertainty quantification by drawing random samples from probability distributions, in which the sampling process does not require knowledge of the entire distribution.

### Prior

The initial probability assigned to a hypothesis before considering new evidence.

### Probability distribution

The statistical description of potential outcomes of random events, where a numerical measure is assigned to the possibility of each specific outcome.

### Probability density estimation

The process of inferring the underlying probability distribution of a random event based on observed data.

### Posterior

The updated probability of a hypothesis after taking into account both prior beliefs and observed evidence.

### Simulation-based inference (SBI)

A statistical method that involves generating synthetic data through forward simulations to make inferences about complex systems, often when analytic or computational solutions are unavailable.

### Virtual brain models (VBMs)

Personalized data-driven models at whole-brain scale that use the connectome for network construction, with a set of equations describing regional brain dynamics placed at each node.

### Information Sharing Statement

All code is provided freely and is available at GitHub (https://github.com/ins-amu/SBI-VBMs) and TVB services on the cloud research platform EBRAINS (https://www.ebrains.eu/).

## Abbreviations

VBM: virtual brain model
SBI: simulation-based inference
SC: structural connectivity
FC: functional connectivity
FCD: functional connectivity dynamic
BOLD: blood-oxygen-level-dependent
fMRI: functional magnetic resonance imaging
ANNs: artificial neural networks
NFs: normalizing flows
MAF: masked autoregressive flow
NSF: neural spline flow
MCMC: Markov chain Monte Carlo

## Acknowledgements

This research has received funding from EU’s Horizon 2020 Framework Programme for Research and Innovation under the Specific Grant AgreementsNo. 945539 (Human Brain Project SGA3), No. 101147319 (EBRAINS 2.0 Project), and No. 101137289 (Virtual Brain Twin Project). The funders had no role in study design, data collection and analysis, decision to publish, or preparation of the manuscript. We thank Jan Fousek, Giovanni Rabuffo, and Paul Triebkorn for fruitful discussions. We acknowledge the use of Fenix Infrastructure resources, which are partially funded from the European Union’s Horizon 2020 research and innovation programme through the ICEI project under the grant agreement No. 800858

## Author contributions

Conceptualization: M.H., M.W, S.P., and V.K.J. Methodology: M.H, A.Z., M.W. Software: M.H., A.Z., and M.W. Investigation: M.H., and A.Z. Visualization: M.H., and A.Z. Supervision: S.P., and V.K.J. Funding acquisition: V.K.J. Writing - original draft: M.H. Writing - review & editing: M.H, A.Z, M.W., S.P., V.K.J.

## 6. Supplementary

**Figure S1.**
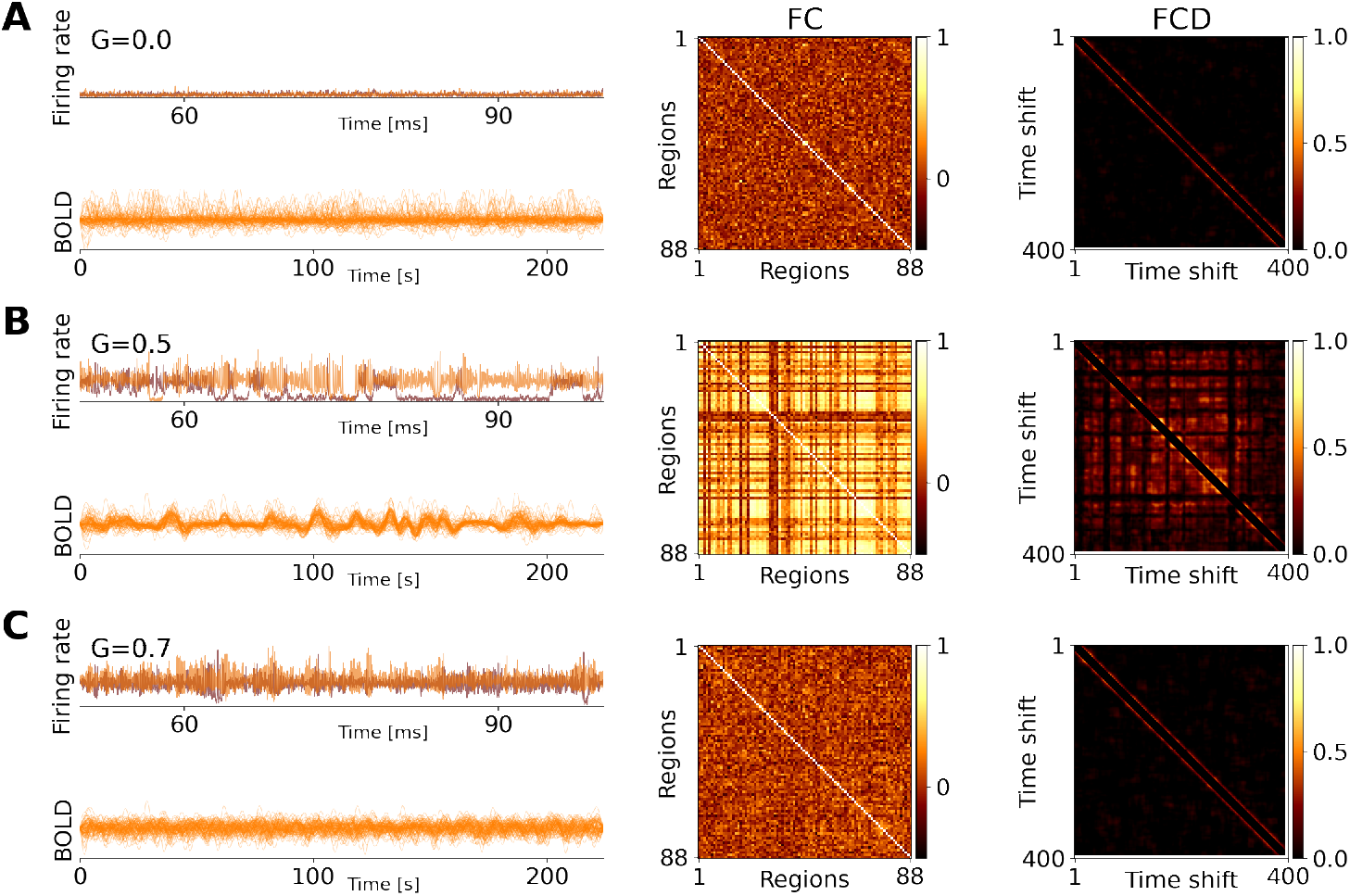
The simulated firing rates and the corresponding BOLD signals across brain regions, for different values of the network scaling parameter G (see Eq. 2), which modulates the overall impact of the SC matrix (left column). The FC (middle column) and FCD (right column) matrices are shown for the simulated BOLD data. (**A**) When the global coupling parameter *G* is weak, the brain regions enter the monostable regime (down-state), resulting in low firing rates across regions. This is manifested as a lack of interactions between different brain regions, resulting in a random FC matrix. Consequently, there are no temporal dynamics in FC, leading to a zero switching index in the FCD matrix (with the variance of 0.001). (**B**) At the working point (here around *G* = 0.5 with noise level of *σ*^2^ = 0.03), the brain regions transition into the bistable regime. This leads to structured transitions between low and high firing rates in those regions. Consequently, the brain demonstrates correlated activities between brain regions, as captured by the FC matrix. Additionally, there is recurrence in the brain’s large-scale dynamics, which is reflected by a non-zero switching index in the FCD matrix (with the variance of 0.009). (**C**) When the global coupling parameter G is strong, the brain regions enter the monostable regime (up-state), resulting in high firing rates across regions. Again, this results in decorrelated regional activities, which are represented by a random FC matrix and a zero switching index in the FCD matrix ((with the variance of 0.002)).

**Table S1.** Sensitivity analysis conducted based on KS and KL distances between the distributions of values in the observed and predicted FC/FCD matrices.

| Hessian value | $G$ | $\alpha$ | $\beta$ |
| --- | --- | --- | --- |
| KS distance between FC matrices | 0.1022 | 0.0085 | 0.0016 |
| KS distance between FCD matrices | 0.2052 | 0.0213 | 0.0033 |
| KL distance between FC matrices | 0.4128 | 0.0337 | 0.0046 |
| KL distance between FCD matrices | 0.42512 | 0.0484 | 0.0109 |

**Figure S2.**
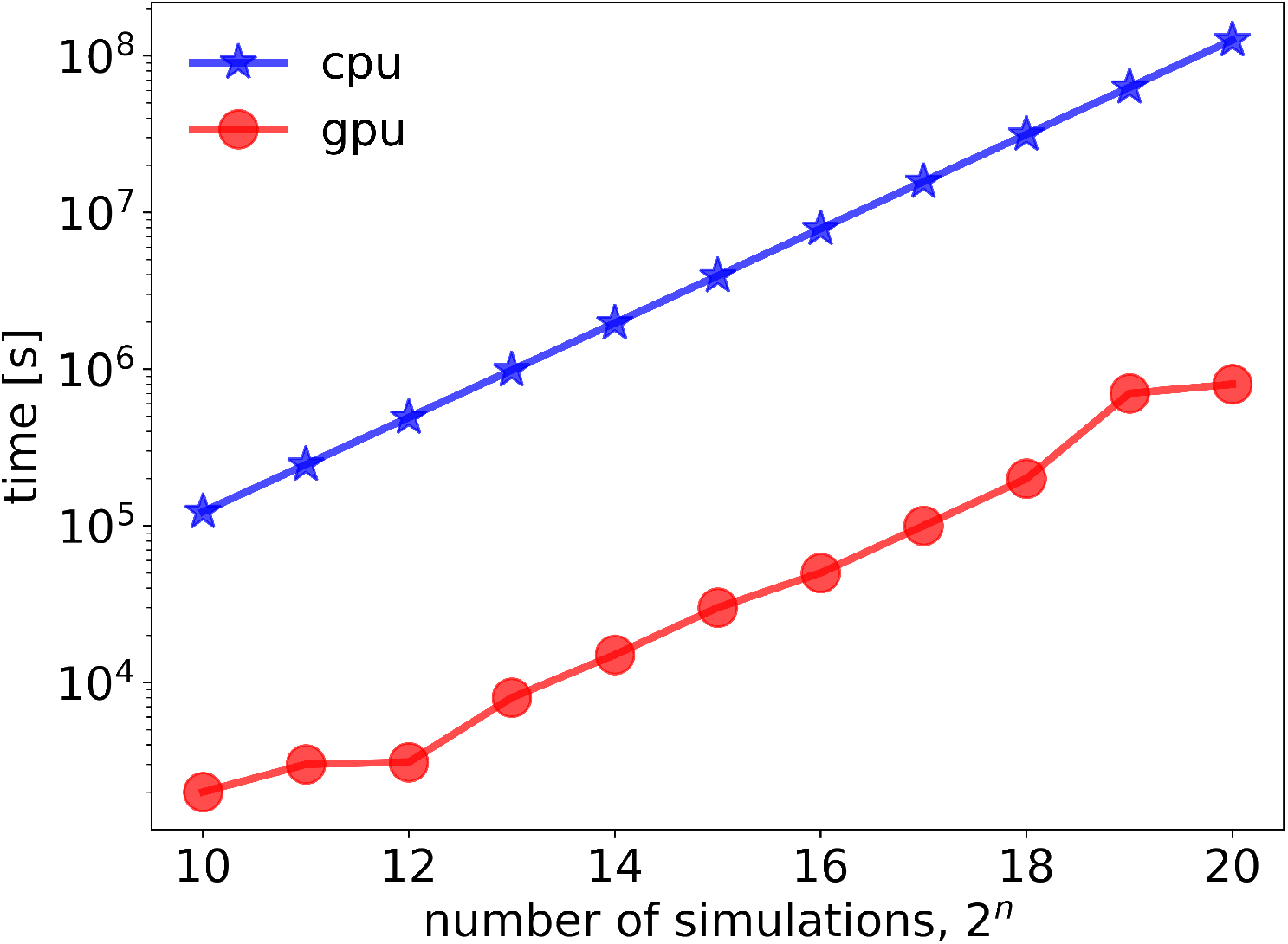
Benchmarking the simulation cost of the VBM using MPR neural mass model, with 2.5M time step on Intel(R) Core(TM) i9-10900 CPU 2.80GHz (in red) and NVIDIA RTX A5000 GPU (in blue). GPUs deliver substantial speedups up to 100X over multi-core CPUs.

**Figure S3.**
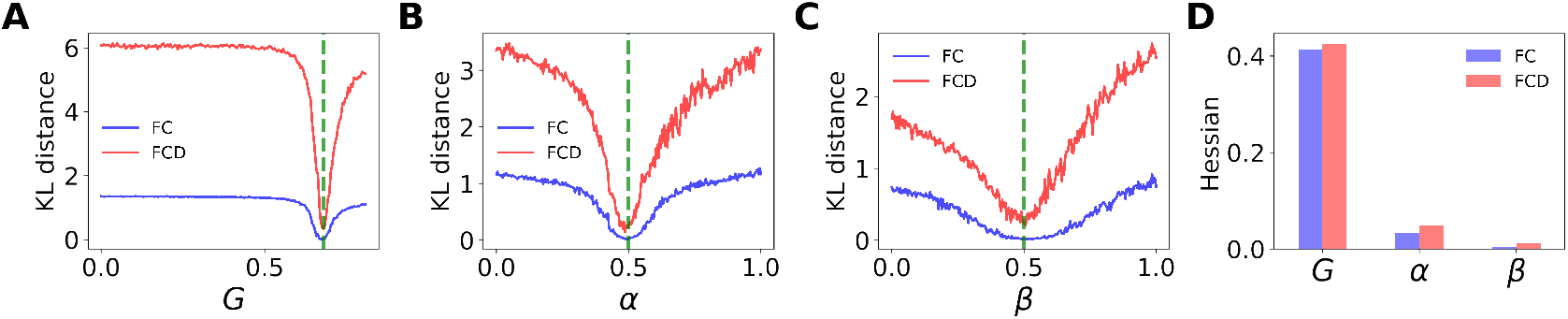
Sensitivity analysis on the degree of degradation in a personalized connectome caused by virtual disorders, based on KL divergence. The plots represent the results of a grid search over the KL divergence between the distribution of observed and predicted FC (in blue) and FCD (in red) values, computed for incremental increase in (**A**) the global scaling parameter *G*, (**B**) the level of deterioration in inter-hemispheric connections *α*, and (**C**) the degree of intra-hemispheric degradation *β* within the limbic system. (**D**) The Hessian values, quantifying the local curvature of KL distance at the ground truth values (in green), using FC and FCD, show no sensitivity to the *β* mask.

**Figure S4.**
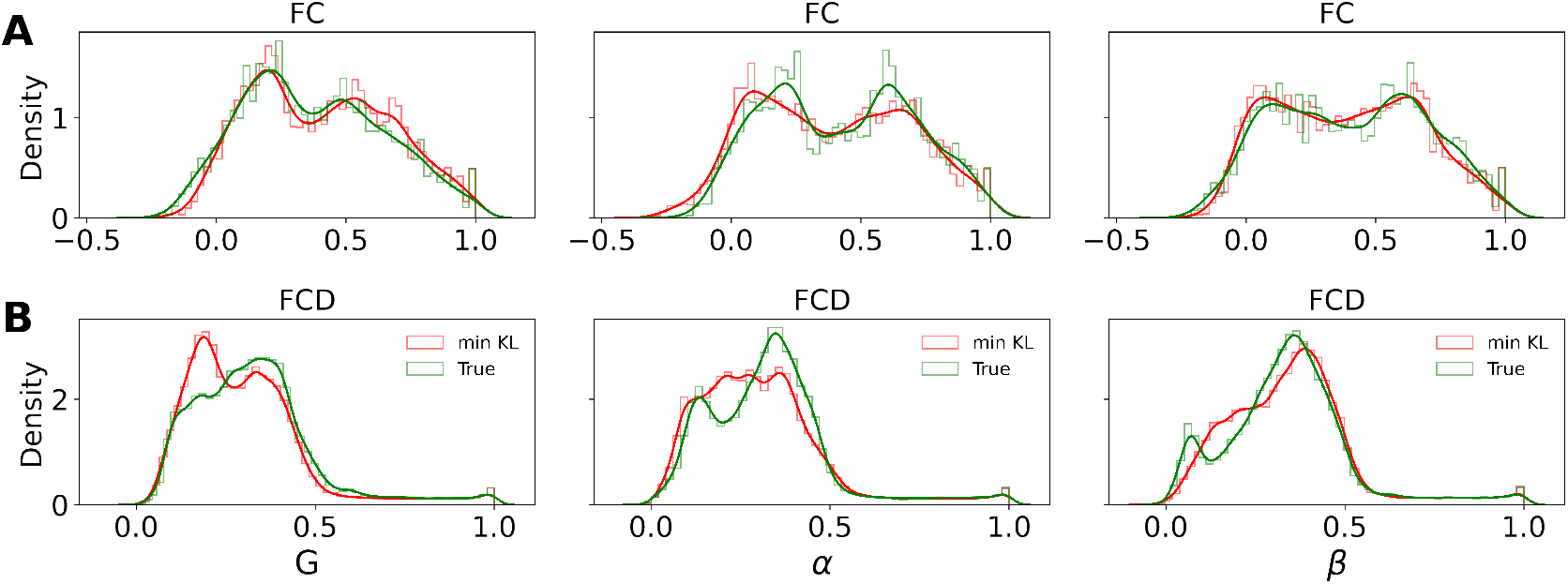
Kullback-Leibler (KL) divergence between the observed and predicted data for estimated parameters over the degree of degradation in the global, between and within connections (denoted by *G, α*, and *β*, respectively) in virtual brains. The estimation is performed by minimizing the KL divergence, which measures the difference between probability distributions. (**A**) and (**B**) display the distribution of FC (functional connectivity) and FCD (functional connectivity dynamics) matrix elements, respectively, for the observed and the estimated working point.

**Figure S5.**
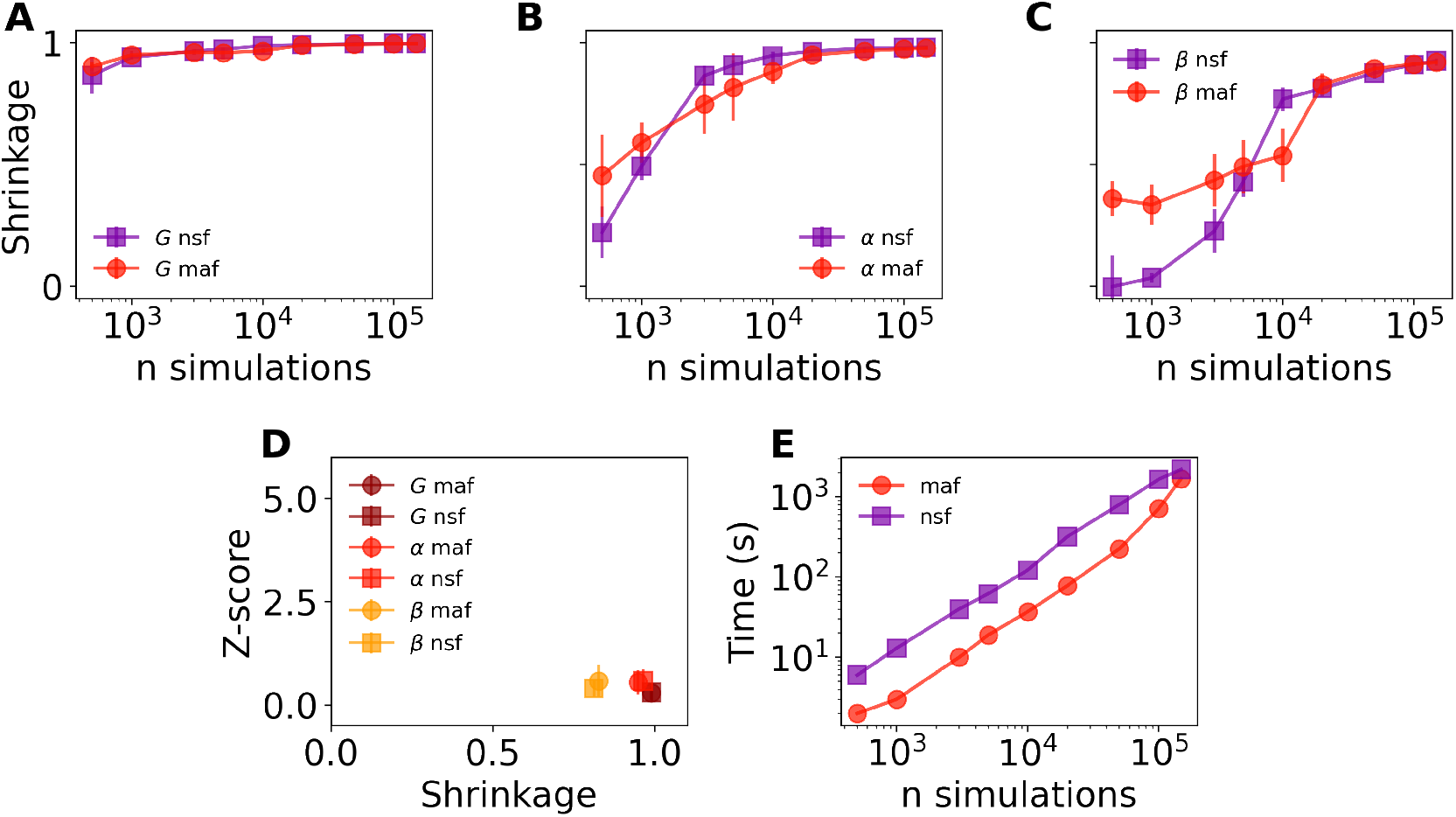
The effect of the number of simulations on uncertainty estimation with training the state-of-the-art deep neural density estimators (MAF and NSF models). (**A**), (**B**), and (**C**) show the plot of posterior shrinkage versus the number of simulations for estimating the parameters *G, α*, and *β*, respectively, from FC/FCD, indicating that both MAF and NSF yield compatible results. (**D**) However, we observed that MAF was 2-4 times faster than NSF during the training process. (**E**) The plot of z-score versus shrinkage indicates an ideal Bayesian estimation for G and α, but not for the β mask.

**Figure S6.**
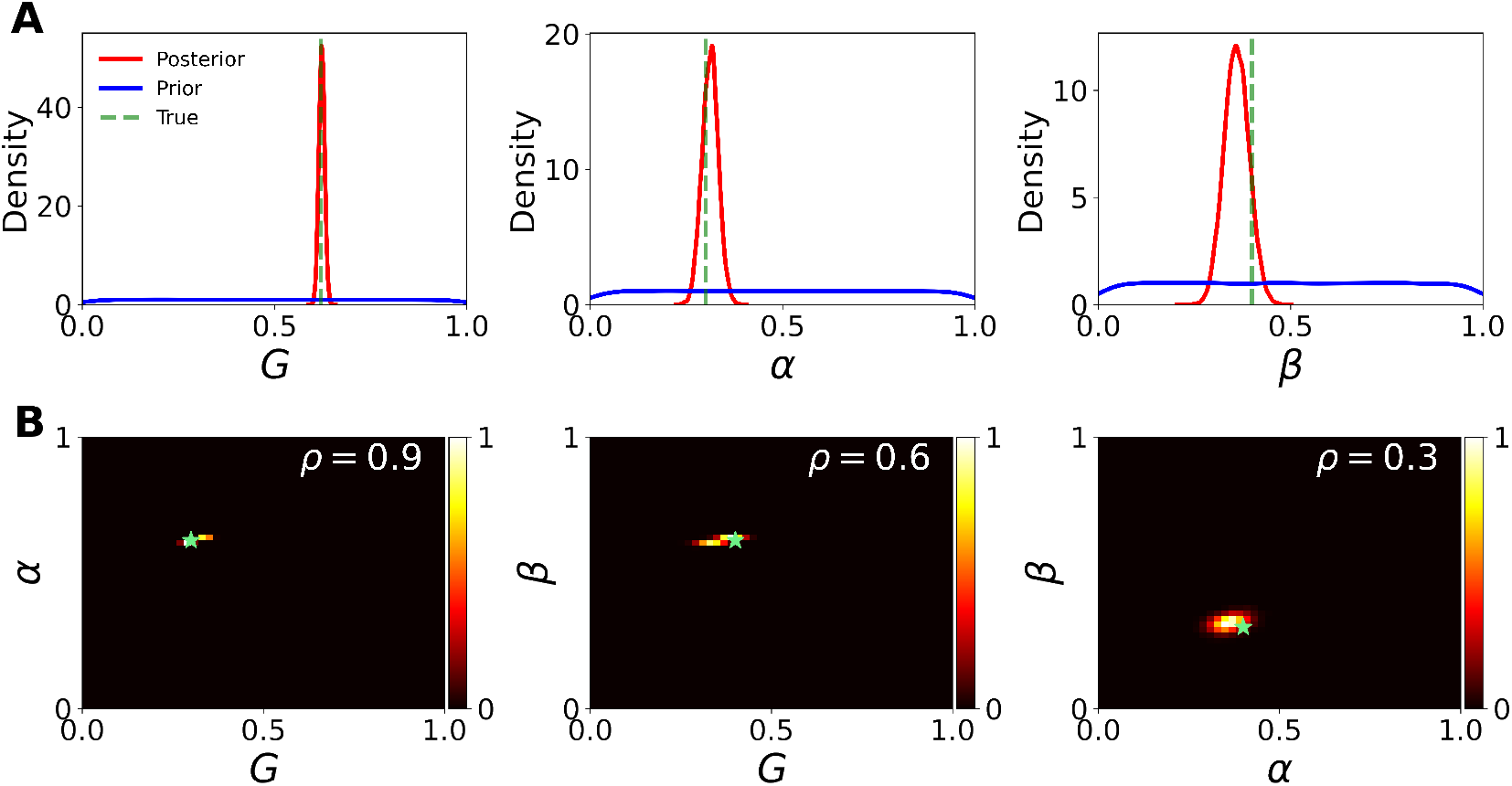
SBI on the global, inter- and intra-hemispheric degradation in a virtual brain, by adding spatio-temporal information to the set of data features. Here we used 100k simulations for training, on both spatio-temporal (statistical moments of the BOLD time-series) and functional (FC/FCD matrices) data freatures. (**A**) The estimated posterior (in red) is shown for the global coupling parameter *G*, the inter-hemispheric degradation mask α, and the intra-hemispheric degradation mask *β*. For random simuations, the parameters are drawn from a uniform prior (in blue). The ground truth values are *G* = 0.62, *α* = 0.3, *β* = 0.4 (in green). We observe that the estimated posteriors accurately capture the true parameters, and they exhibit appropriate shrinkages from the priors. (**B**) The joint posterior distributions indicate a correlation of *ρ* = 0.9 between *G* and *α*, a correlation of *ρ* = 0.6 between *G* and *β*, and a correlation of *ρ* = 0.3 between *α* and *β*.

**Figure S7.**
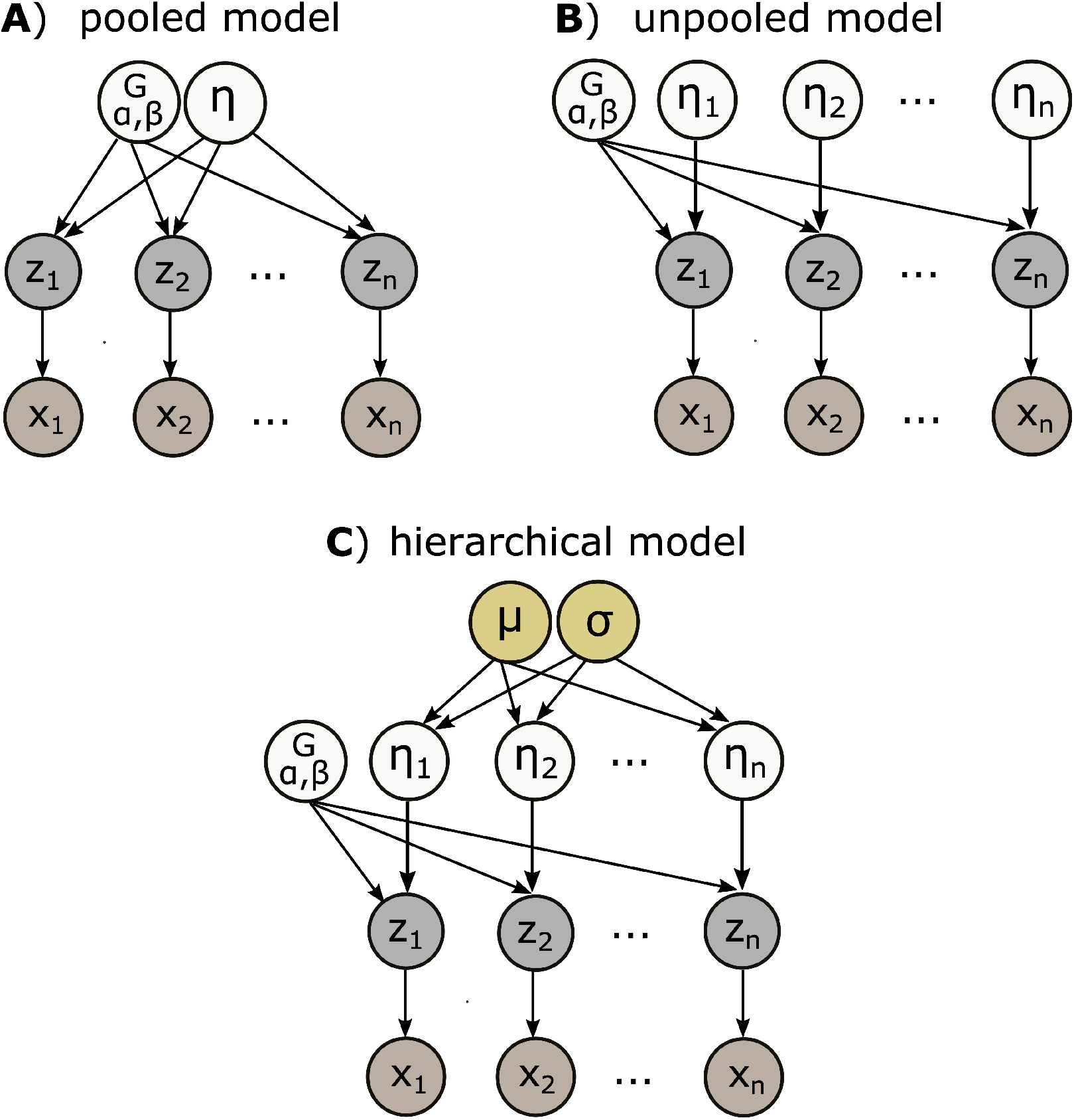
Modeling structures in Bayesian framework, to explains the observation denoted by *xi* (e.g., BOLD data) through the hidden layer of *z*_*i*_ (e.g., firing rates in Eq. (2)). (**A**) Pooled (homogeneous) modeling, which assumes a common distribution across all individual parameters and assumes no variation in the sampling process. Hence, the set of generative parameters is low-dimensional as 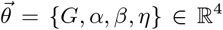. (**B**) Unpooled (heterogeneous) modeling captures the distinct characteristics of individual parameters without imposing a uniform structure. Hence, the set of generative parameters becomes high-dimensional as 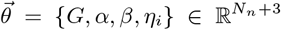 with i *∈ {*1, 2, …, *N*_*n*_}. (**C**) Hierarchical (or partial pooling) modeling offer a balance between the simplicity of pooled models and the complexity of unpooled models. Hence, the set of generative parameters is optimal as 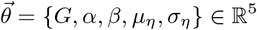,with 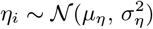.

**Figure S8.**
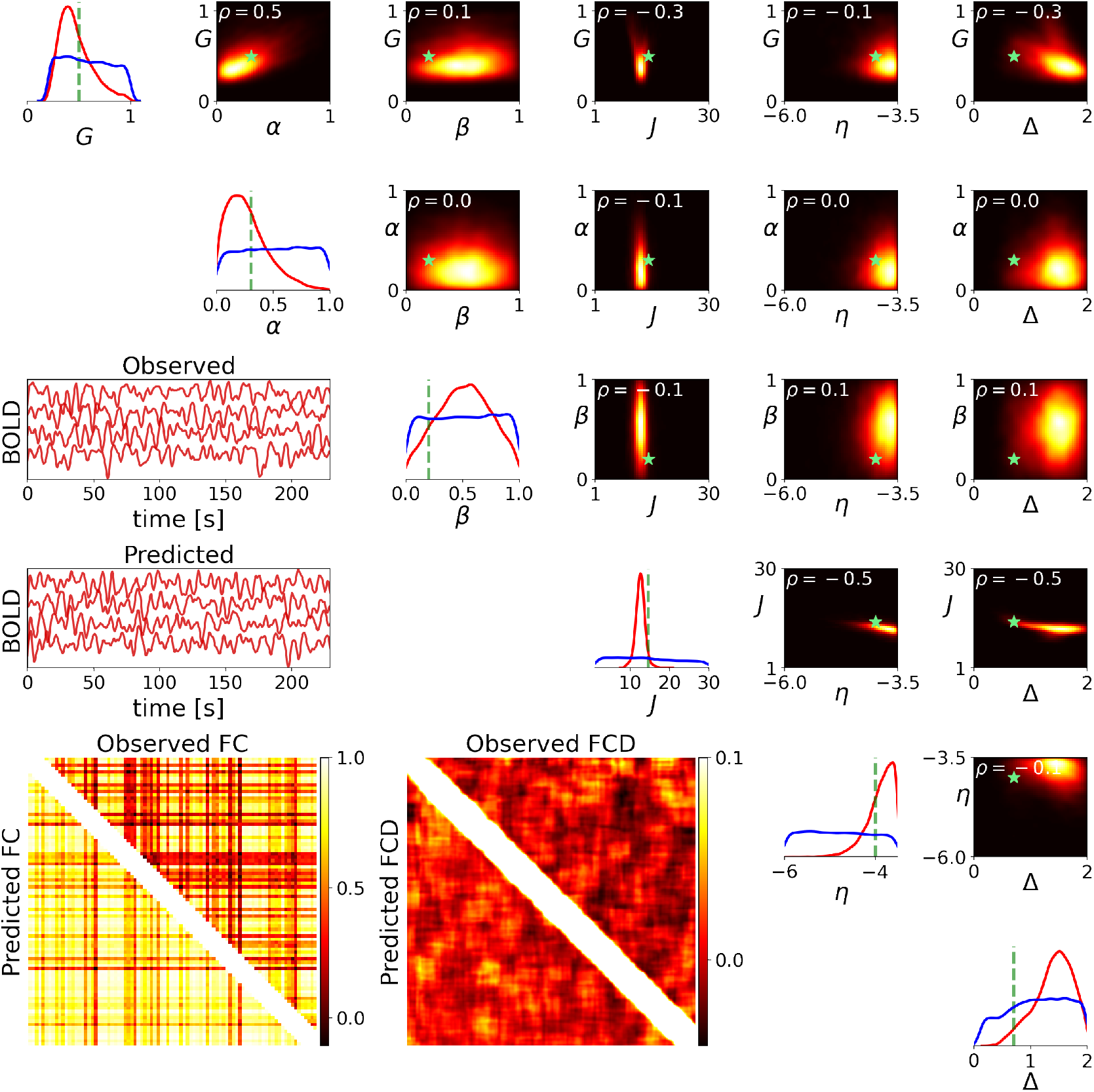
Inference on homogeneous generative parameters in a virtual brain using only functional data features. Here the set of inferred parameters is 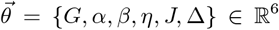. The training process involved using only functional data features (FC/FCD), with a budget of 100k simulations. The diagonal panels display the prior (in blue) and estimated posterior (in red). For most of parameters, the maximum a posterior deviates form true value (green lines) with a large uncertainty. The lower diagonal panels display the observed and predicted BOLD data and the corresponding FC/FCD matrices. The upper diagonal panels show the joint posterior between parameters, and their correlation values (*ρ* at the upper left corners). The ground-truth values are shown by green stars. High-probability areas are color-coded in yellow, while low-probability areas are represented in black.

**Figure S9.**
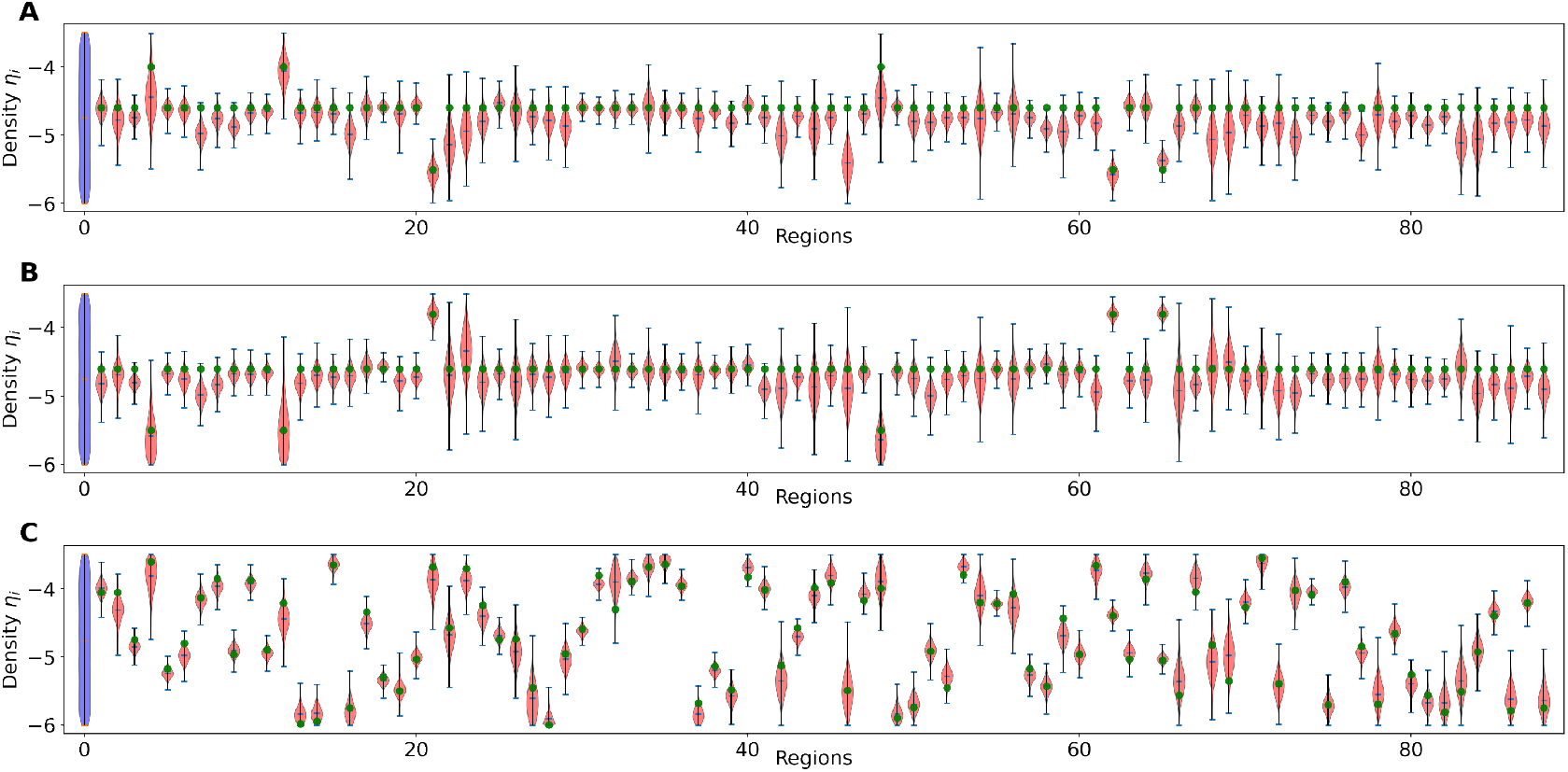
Inference on heterogeneous excitabilities *η*_*i*_ for 88 brain regions. SBI provides accurate posterior estimation (in red) using both spatio-temporal and functional data features for (**A**)-(**C**) different configurations in excitabilities across brain regions (ground truth in green dots). Here, we used 1M random simulations from uniform priors (in blue) as: *η*_*i*_ ∈ 𝒰 (*−*6, *−*3.5).

**Figure S10.**
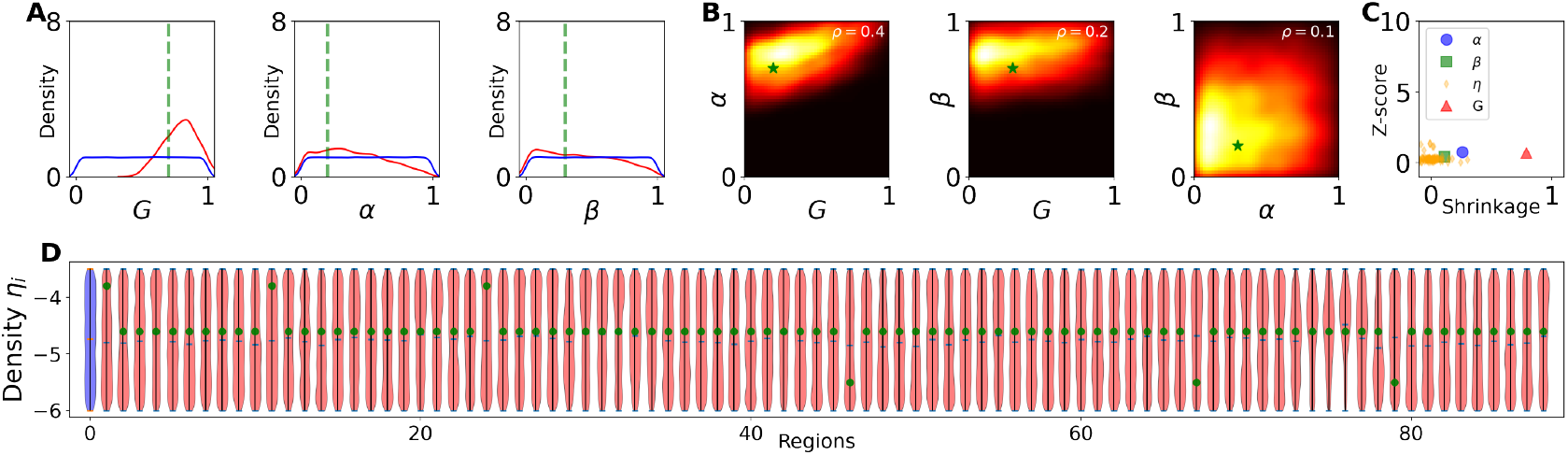
SBI on the set of generative parameters denoted by 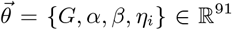, where *η*_*i*_ with *i ∈ {*1, 2, …, *N*_*n*_ = 88*}*. The training process involved only functional data features, on a budget of 1M simulations. (**A**) The estimated posterior (in red) for parameters *G, α*, and *β*. The prior (in blue), for each was placed between zero and one. The ground truth values are *G* = 0.62, *α* = 0.3, *β* = 0.4 (in green). (**B**) The join posterior between parameters. (**C**) The plot of z-score versus shrinkage indicates an ideal estimation for *G* and *α*, but poor estimation for *β* and *η*_*i*_. (**D**) The estimated posterior of *η*_*i*_, for 88 regions. The prior on excitabilities (in blue) was placed as: *η*_*i*_ ∈ 𝒰 (*−*6, *−*3.5). This result indicate that relying on functional data is inadequate for accurate inference on heterogeneous excitabilities distributed across the whole brain regions.

**Figure S11.**
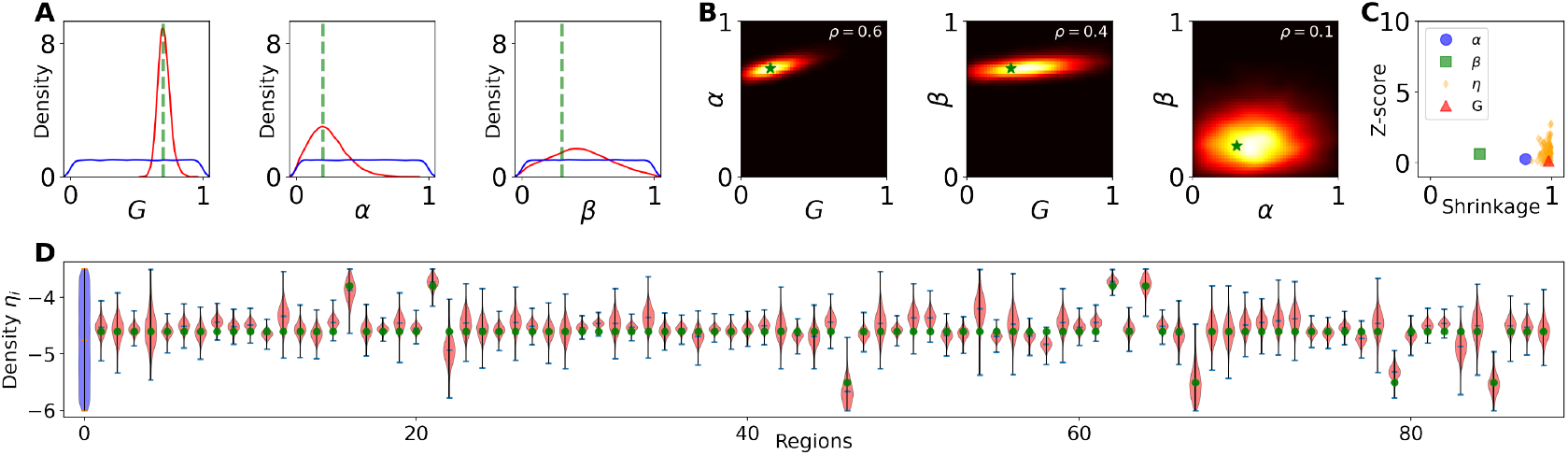
Estimation on heterogeneous generative parameters, using MAF model for training in SBI. Here the set of generative parameters is 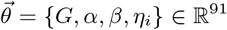, with *i ∈ {*1, 2, …, *N*_*n*_ = 88*}*. The training process involved using both spatio-temporal and functional data features, with a budget of 1M simulations. (**A**) The estimated posterior (in red) for parameters *G, α*, and *β*. The prior (in blue), for these parameters was placed between [0,1]. The ground truth values are *G* = 0.62, *α* = 0.3, *β* = 0.4 (in green). (**B**) The join posterior between parameters. (**C**) The plot of z-score versus shrinkage indicates an ideal estimation for *G, α*, and *η*_*i*_ but poor estimation for *β*. (**D**) The estimated posteriors of *η*_*i*_, for 88 regions (in red) demonstrate no shrinkages from prior (in blue). The prior on excitabilities was placed as: *η*_*i*_ ∈ 𝒰 (*−*6, *−*3.5). This result indicates that both spatio-temporal and functional data features provides an ideal Bayesian estimation on all generative parameters, except the intra-hemispheric degradation within the limbic system denoted by *β* mask.

**Figure S12.**
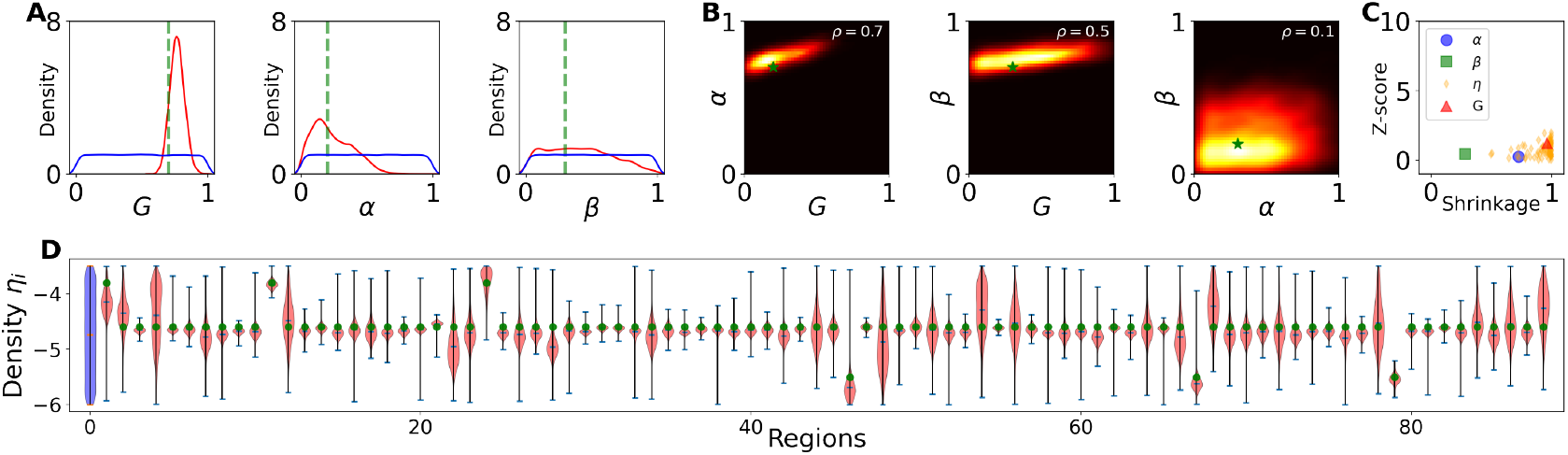
The same analysis as shown in Fig S11, but using the NSF model for training in SBI. Both the MAF and NSF models are compatible in accuracy of estimation and uncertainty quantification of the generative parameters.

**Figure S13.**
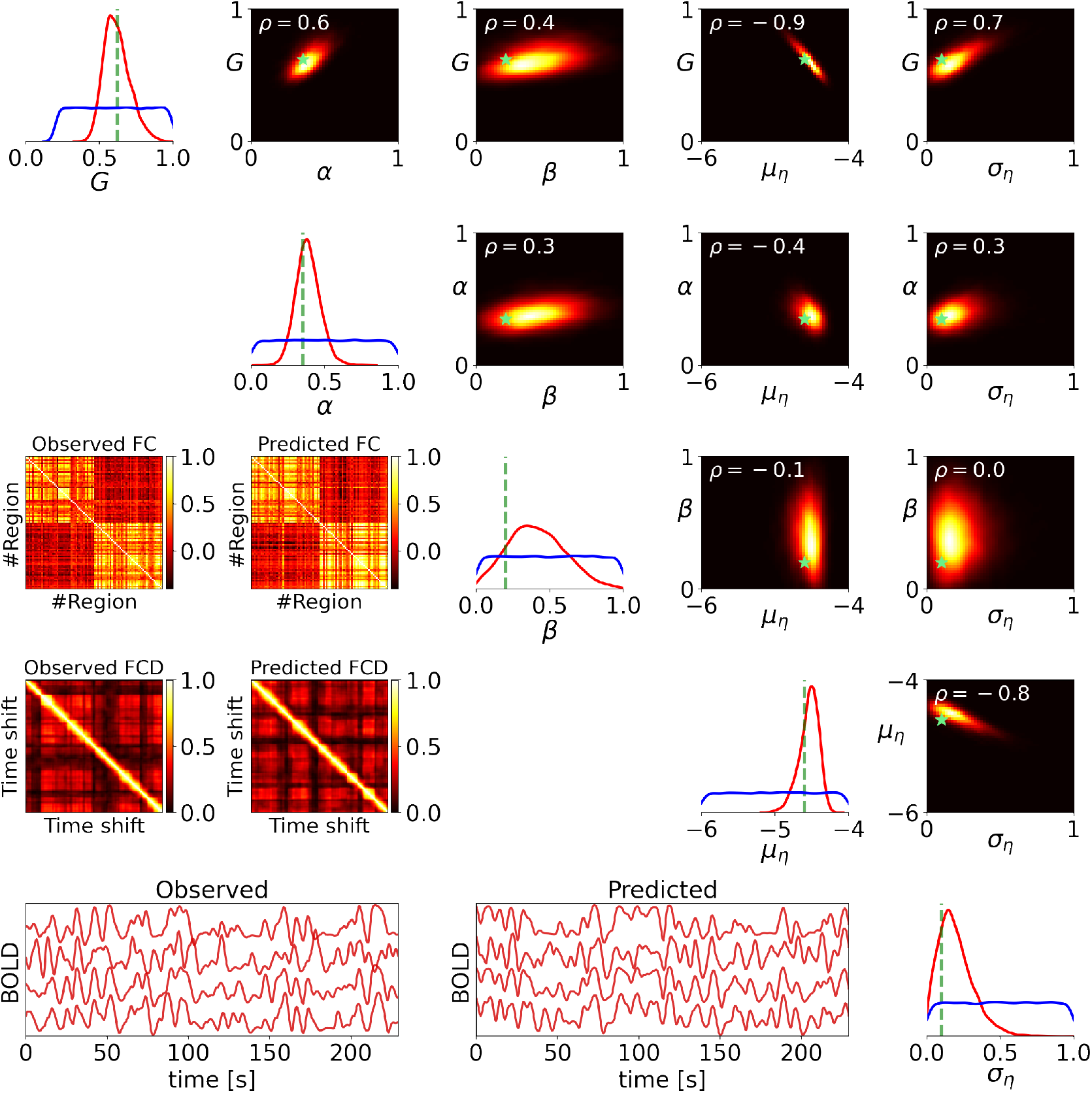
The same analysis as shown in Fig 5, but excluding the spatio-temporal data feature from training. In contrast to heterogeneous modeling (Figure S9), the hierarchical modeling provides informative posteriors when using only functional data feature for inference, even with a significantly smaller budget of simulations.

